# Cardiac differentiation roadmap for analysis of plasticity and balanced lineage commitment

**DOI:** 10.1101/2024.04.09.588683

**Authors:** Rebecca R. Snabel, Carla Cofiño-Fabrés, Marijke Baltissen, Verena Schwach, Robert Passier, Gert Jan C. Veenstra

## Abstract

Stem cell-based models of human heart tissue and cardiac differentiation employ monolayer and 3D organoid cultures with different properties, cell type composition, and maturity. Here we show how cardiac monolayer, embryoid body, and engineered heart tissue trajectories compare in a single-cell roadmap of atrial and ventricular differentiation conditions. Using a multiomic approach and gene-regulatory network inference, we identified regulators of the epicardial, atrial and ventricular cardiomyocyte lineages. We identified *ZNF711* as a regulatory switch and safeguard for cardiomyocyte commitment. We show that *ZNF711* ablation prevents cardiomyocyte differentiation in the absence of retinoic acid, causing progenitors to be diverted more prominently to epicardial and other lineages. Retinoic acid rescues this shift in lineage commitment and promotes atrial cardiomyocyte differentiation by regulation of shared and complementary target genes, showing an interplay between *ZNF711* and retinoic acid in cardiac lineage commitment.

**GRAPHICAL ABSTRACT:** 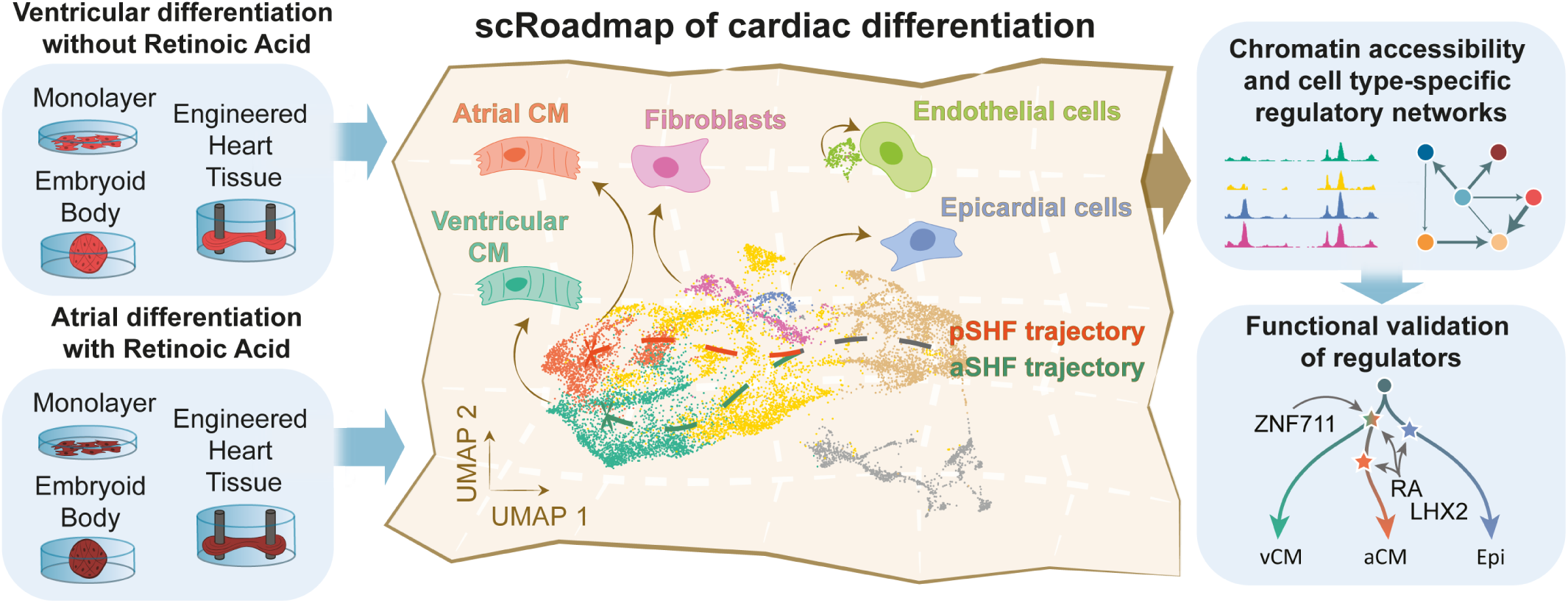

**HIGHLIGHTS:**

- We present an atlas of cardiac differentiation under different culture conditions, showing progenitor trajectories and cell states reminiscent of the fetal heart.
- Multiome and chromatin accessibility analysis identified candidate regulators of cardiomyocyte and non-cardiomyocyte lineages.
- Distinct trajectories of lineage commitment and differentiation under different culture conditions shows plasticity of first and second heart field progenitors and different ratios of cardiomyocyte and non-myocyte lineages.
- ZNF711 is required for balanced commitment to epicardial and ventricular cardiomyocyte lineages. Retinoic acid rescues a requirement for ZNF711 in cardiomyocyte differentiation, promoting both atrial cardiomyocyte and epicardial differentiation.

## INTRODUCTION

The first precursor cells of the heart emerge during early gastrulation when the three germ layers of the embryo are formed (Lescroart et al., 2014; Saga et al., 1999). These progenitor cells reside within the splanchnic lateral mesoderm in spatially segregated populations, notably the first and the second heart field (FHF and SHF), which together will form the cardiac crescent (Buckingham et al., 2005). These progenitor states and their lineage contributions have been studied using single-cell omics approaches in mouse embryonic tissues, as reviewed recently (Lescroart & Zaffran, 2022; Sendra et al., 2021; Tyser, 2023). The lineages of the FHF primarily populate the left ventricle and part of the atria. The SHF is divided into an anterior and posterior section (aSHF and pSHF), which are transcriptionally distinct and contribute to different parts of the developing heart (Lescroart et al., 2018). The aSHF forms the right ventricular myocardium and outflow tract of the heart, whereas progenitors of the pSHF commit to the atrial lineage under the influence of retinoic acid (RA) signaling (Protze et al., 2019). Reports on cardiac development in humans start at five weeks of gestation, when the heart already features multiple chambers (Asp et al., 2019; Cui et al., 2019; Devalla et al., 2015; Farah et al., 2024; Sahara et al., 2019). By that time, the multipotent cardiac progenitors have produced the cardiomyocyte, endothelial, epicardial and smooth muscle lineages and have built a myocardium that forms the atria and ventricles (Protze et al., 2019). Very little is known about the earliest stages and distinctive lineage choices of human cardiac development *in vivo* (Tyser et al., 2021; Zeng et al., 2023).

Human pluripotent stem cells (hPSC) offer an immense potential to differentiate many cell types *in vitro*, thus presenting a relevant alternative in mimicking the earlier steps of human tissue development (Takahashi et al., 2007; Thomson et al., 1998; Yu et al., 2007). Various culture models have been established to obtain cardiomyocytes (CMs) (Burridge et al., 2012; Devalla & Passier, 2018; Mummery et al., 2012), with variations including an early step of retinoic acid (RA) treatment to enrich for atrial CMs (aCMs) (Devalla et al., 2015; Lee et al., 2017; Schwach et al., 2022; Zhang et al., 2010) or epicardial derivatives (Guadix et al., 2017; Hofbauer et al., 2021a; Meier et al., 2023). Over time, the cardiac cultures have increased in complexity (e.g. from two- to three-dimensional cultures and combinations of different cell types) leading to advanced maturity of the cardiomyocytes and simultaneous development of a dynamic pool of cardiac cell types (DeLaughter et al., 2016; Kannan et al., 2021).

Single-cell transcriptomic studies have elucidated the differentiation processes of hPSCs towards CMs (Ameen et al., 2022; Churko et al., 2018; Friedman et al., 2018; Grancharova et al., 2021; Hofbauer et al., 2021b; Lewis-Israeli et al., 2021; Meier et al., 2023; Mononen et al., 2020; Ruan et al., 2019; Sahara et al., 2019; Zawada et al., 2023). Several of these studies highlighted the presence of non-myocyte subpopulations within the culture (Churko et al., 2018; Friedman et al., 2018; Mononen et al., 2020), while others shed light on the specification of particular cardiomyocyte subtypes, such as atrial, right and left ventricular CMs, and node-like CMs (Funakoshi et al., 2021; Pezhouman et al., 2021; Wiesinger et al., 2022; Yang et al., 2022). However, very little is known about how cardiac multipotent progenitors give rise to different cardiomyocyte and non-myocyte lineages, and how this is regulated. Hence, there is a need for disentangling the gene regulatory networks (GRNs) of cardiac development as the cells commit to specific lineages, and how they govern cardiac differentiation in the context of *in vitro* culture conditions. Understanding these mechanisms will aid in the advancement of cardiac hPSC models and their ability to more accurately model cardiovascular development and heart disease.

Here we present a detailed comparison of *in vitro* cardiac differentiation conditions from hPSCs using single-cell transcriptomic (RNA) and chromatin accessibility (ATAC) data, which we used to construct a roadmap of atrial and ventricular multilineage differentiation. We used a combination of monolayer, embryoid body (EB) and engineered heart tissue (EHT) cultures to compare their cellular compositions and differentiation stages. In addition, we used RA to compare multilineage differentiation in atrial and ventricular cultures. Direct comparisons between the cardiac models revealed subpopulations resembling *in vivo* cell states and identified transcriptional regulators at the onset of lineage commitment. The integrative multiomic approach led to the identification of *ZNF711*, and the interplay of this zinc finger with RA in balancing commitment to the cardiomyocyte and epicardial lineages.

## RESULTS

### Cardiomyocyte differentiation methods produce cardiac cell states found in the human fetal heart

To compare the early steps of cardiac lineage development, we differentiated dual fluorescent *NKX2-5^EGFP/+^*and *COUP-TFII* (*NR2F2*)*^mCherry/+^* hPSCs (Schwach et al., 2017) to CMs under cardiomyocyte monolayer (Mon) and cardiac EB conditions, with or without RA (*see Methods*, Fig. 1A, Supplementary Fig. S1A). We and others showed previously that RA redirects differentiation towards aCMs (Devalla et al., 2015; Schwach et al., 2017, 2022; Zhang et al., 2010), which occurs at high efficiency as evaluated by fluorescence activated cell sorting (Supplementary Fig. S1B). At day 14, CMs of these cultures were further matured for an additional 12 days (m12) until day 26, by either generating 3D engineered heart tissues (EHT) (Birket et al., 2015; Ribeiro et al., 2022), or replating to monolayer as a control for the changes that occur in the EHTs, both with the addition of cardiomyocyte maturation medium (Supplementary Fig. S1C). The atrial and ventricular CMs expressed the expected regional markers (NR2F2 and MYL2 respectively; Fig. 1B) in addition to general cardiomyocyte markers (MYL7, ACTN2; Supplementary Fig. S1D), while displaying typical action potential profiles, with higher frequencies and shorter action potential duration times in aCMs (Supplementary Fig. S1E). Continued culture of the CMs in the EHT model enhanced their respective identities, as shown by immunohistochemistry of aCM- and vCM-specific markers (Supplementary Fig. S1F).

**Figure 1.**
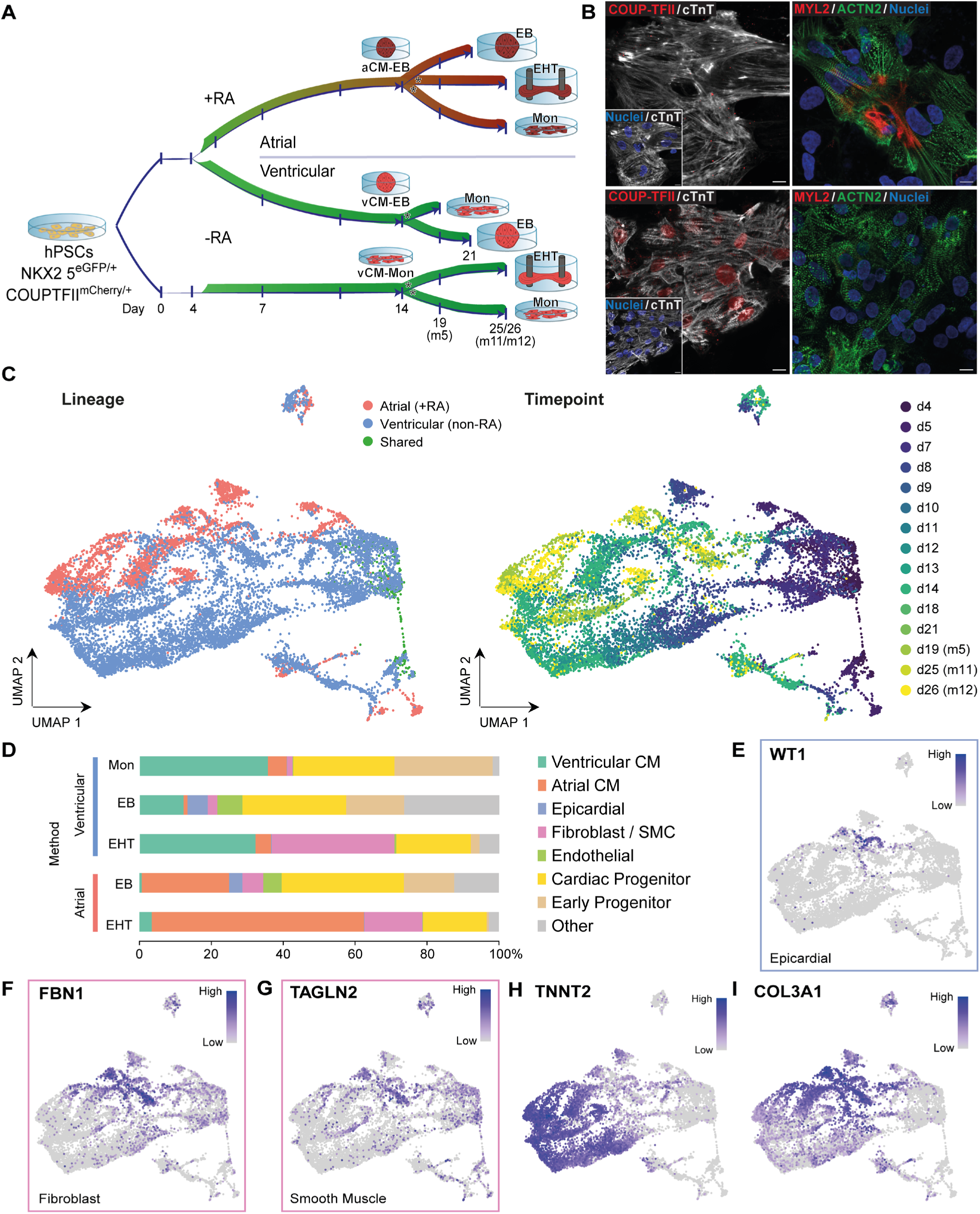
Confirming cardiac cell types *in vitro* with *in vivo* comparison. (A) Human pluripotent stem cell (hPSC)-derived cardiomyocyte differentiation cultures set up. hPSCs are differentiated in monolayer (Mon) or embryoid body (EB) cultures over the course of 14 days. At day 14, sorted CMs and non-CMs are replated in engineered heart tissues (EHT) to further mature, or replated as monolayers for comparison of the 3D culture (indicated with stars), while the EBs are kept in culture until day 21 (detailed overview of differentiation protocols in Supplementary Fig. S1). RA = retinoic acid, CM = cardiomyocyte, Mon = monolayer, EB = embryoid body, EHT = engineered heart tissue. d1-21 = day 1-21, m5-m12 = day 5-12 in maturation medium (day 14-26; EHT and monolayer only). (B) Immunohistochemistry of atrial (COUP-TFII / NR2F2), ventricular (MYL2) and general cardiomyocyte (cTnT / TNNT2 and ACTN2) marker genes in day 14 cardiomyocytes of both the vCM (top row) and aCM (retinoic acid treated; bottom row) culture conditions. Scale bar = 10 (COUP-TFII/cTnT) and 20 μm. (C) Single-cell temporal atlas UMAP labelled with culture conditions (left) and time point of harvest (right). The resulting dataset consists of 2 independent differentiation experiments per culture protocol. (D) Proportions of the different cell states per culture method. Cell states annotation process is shown in Fig. S4. (E-G) Cell type marker expression levels visualized onto the atlas UMAP, with *WT1* as epicardial, *FBN1* as fibroblast and *TAGLN2* as smooth muscle marker. (H-I) Inversely related expression patterns of *TNNT2* (F) and *COL3A1* (G) visualized onto the atlas UMAP.

To produce a detailed map of how cardiac progenitors give rise to cardiomyocytes and other cardiac cell types, we performed single-cell RNA sequencing on samples of different culture methods and time points (Fig. 1A, C and Supplementary Fig. S1A, C). We used a 384-well plate-based platform (SORT-Seq) because of the flexibility it offers in experimental design and it facilitates the capture of many time points and different culture conditions (Hashimshony et al., 2016; Muraro et al., 2016). Particularly, we captured cells at the mesodermal stage (day 4-5), cardiac progenitors (day 7-9), cells committed to the CM fate (day 10-14) and definitive CMs (day 18-26). This resulted in a single-cell landscape covering different protocols and time points with a total of 12,706 cells after filtering for quality (*see Methods*, Supplementary Fig. S2A-B). Known cardiac markers for different stages of development were evaluated for their overall expression patterns over time (Supplementary Fig. S2C), indicating a decrease in mesodermal markers (*MIXL1*, *PDGFRA*), alongside an increase in cardiac progenitor markers (*MEF2C*, *NKX2-5*), followed by structural CM genes (*TNNT2*, *MYL7*) in combination with specific atrial (*KCNA5*, *KCNJ3*, *VSNL1*) or ventricular (*MYL2*, *MYH7*, *VCAM1*) CM genes (Supplementary Fig. S2C-D).

To predict which cells of the different culture methods represent a similar biological state, we integrated the data using an unsupervised selection of mutual cell pairings across the multiple culture experiments (Supplementary Fig. S2E-F) (Haghverdi et al., 2018; Stuart et al., 2019). This resulted in an integrated *in vitro* cardiac differentiation atlas or roadmap, in which a chronological progression of time points and cell states was observed, while resolving distinctions between the atrial and ventricular lineage (Fig. 1C). Comparing the end states of the differentiation culture methods revealed a clear shift of the definitive CM populations between monolayer and EHT cultures, corresponding to further maturation upon culturing as 3D EHTs (Supplementary Fig. S2G). To annotate the different cell states across the different cultures and time points, we compared the cell clusters with an integrated dataset of human fetal heart development (Asp et al., 2019; Cui et al., 2019) (Supplementary Fig. S3A-C and S4A-D). This *in vivo* comparison revealed that from day 8 onward and specifically in the EB method, clusters matching cardiac fibroblasts and smooth muscle cells were present (clusters 15-16), alongside clusters comparable to endothelial and epicardial cells (respectively clusters 17 and 22) (Fig. 1D, Supplementary Fig. S4A-D). Fibroblasts (cluster 16) were relatively abundant in the EHT model, but epicardial-like cells (cluster 22) were not (Fig. 1D, Supplementary Fig. S4B-D). This decrease is likely linked to the dissociation of the EBs and sorting and selection of the cells for the EHTs with an 80:20 ratio of CM and non-CM cells (Supplementary Fig. S1C). Expression of well-known marker genes, such as *FBN1* for fibroblasts, *WT1* for epicardium and *TAGLN2* for smooth muscle cells (Fig. 1E-G), further confirmed the presence of these non-myocyte cell types. Lastly, one set of clusters (clusters 10, 13, 18 and 25) of cells present in the EB cultures of both lineages (Supplementary Fig. S4D), was found enriched for markers of endoderm-committed cell types, such as early foregut progenitors (Supplementary Fig. S5) (Ang et al., 2018; Scheibner et al., 2021). The definitive endoderm and foregut derivatives can contribute to epicardial and myocardial lineage development (Branco et al., 2022). Overall, these data highlight the broad lineage potential in both ventricular and atrial EB cultures (Fig. 1D). In addition, we found that a population of cells consisting of non-CM and early CM progenitors, were marked by high expression of *COL3A1*, and that this expression is inversely related to that of *TNNT2* in the CM lineage (Fig. 1H-I). This *COL3A1*-high population shows a different abundance in the cardiac cultures, raising the possibility that the balance of lineage commitment of multipotent progenitors towards CM and non-CM identities is affected by the culture method.

### Early trajectories show expression patterns mimicking *in vivo* development

To study the relationship between *COL3A1*-high progenitors and the CM and non-CM lineages, we performed cell trajectory analysis for each culture method from early time points up to day 14 (Fig. 2A-F). This revealed the progression of cardiac progenitors expressing *COL3A1* (Fig. 2G-I) to epicardial cells in atrial and ventricular EBs (Fig. 2D-F, Supplementary Fig. S6) and *TNNT2*-positive CMs in all cultures (Fig. 2J-L). The epicardial trajectory is supported by the presence of cells expressing *ISL1, HAND1* and *BNC2*, markers of the juxta-cardiac field or pre-epicardial progenitor state (Supplementary Fig. S6A-D). This population continued to develop into cells negative for *ISL1* and *NKX2-5*, while expressing epicardial markers *BNC2*, *WT1* and *ITGA8* (Supplementary Figs. S6B, E-G). In each of the early cultures, the *COL3A1*-high state was present, although this was more pronounced in the atrial EBs treated with RA (Fig. 2I). In the ventricular monolayer culture, CM progenitors can go through either a high or a low *COL3A1*-expression state before acquiring the CM identity (Fig. 2G, J). In EBs, the progenitors are routed almost exclusively through a *COL3A1*-high state (Fig. 2H-I), but the *COL3A1*-high population is much expanded in the RA-treated atrial EB culture conditions. In most progenitors present in the EBs, *COL3A1* appears to be co-expressed with the second heart field (SHF) progenitor marker *ISL1* (Fig. 2H-I, Supplementary Fig. S6B), as shown before (Mononen et al., 2020). There is no co-expression, however, in the epicardial population, which does express *COL3A1* but not *ISL1* (Fig. 2H-I, Supplementary Fig. S6B). This is in line with previously described dynamics of *ISL1* expression during epicardial lineage progression (Meier et al., 2023).

**Figure 2.**
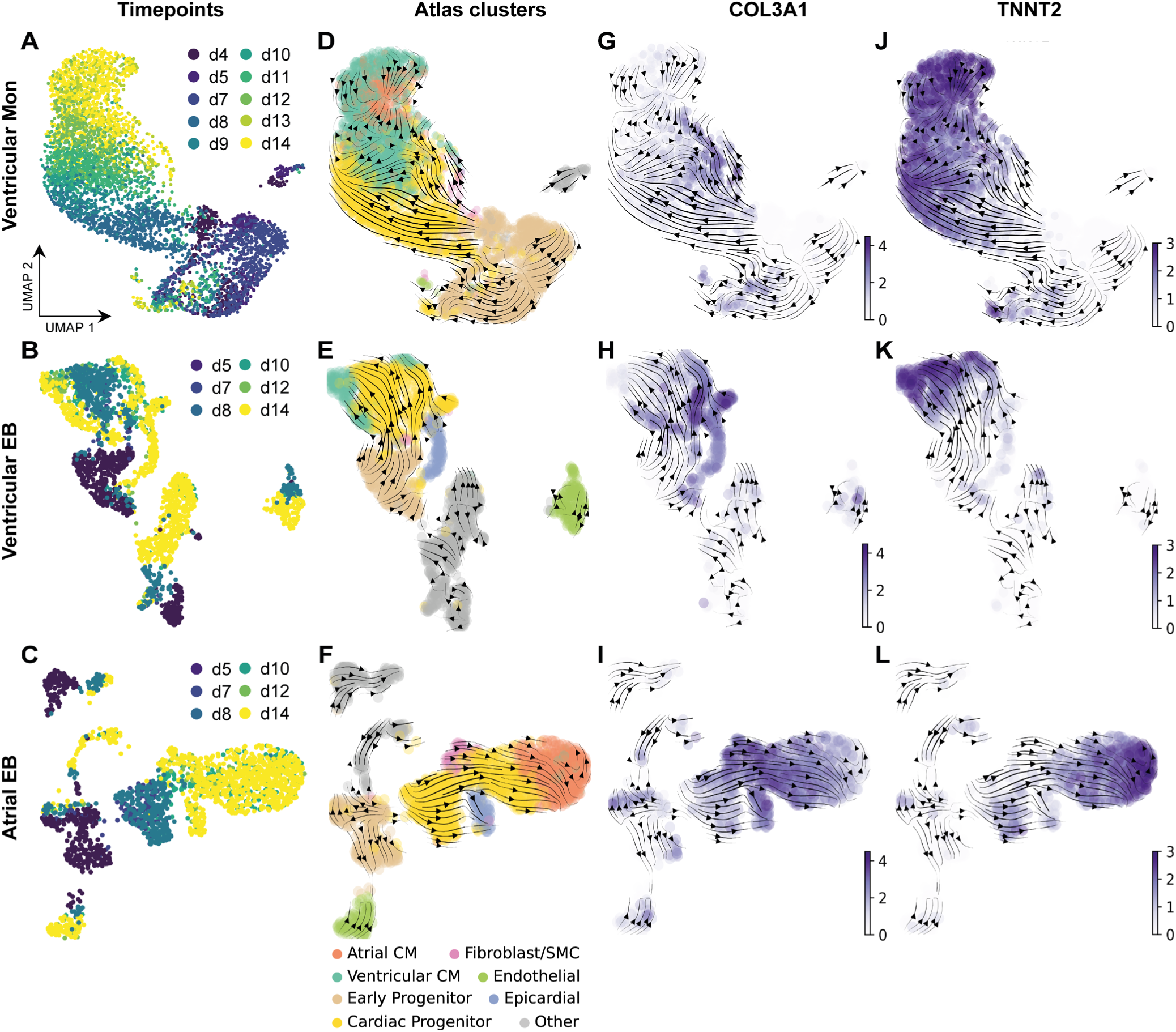
Cardiac progenitor differentiation trajectories in EB and monolayer cultures. UMAP representations of early time point (day 4 to day 14) subsets of the atlas per culture protocol (top, middle and bottom row representing respectively the vCM monolayer, ventricular EB, and atrial EB protocols). (A-C) UMAPs labelled with time points (d4-d14, days of differentiation). (D-F) UMAPs labelled with atlas cell states (cf. Fig. 1D and Supplementary Fig. S4C). (G-I) and (J-L) UMAPs with the expression values of respectively *COL3A1* and *TNNT2*. Panels (B-L) additionally show arrows for cell state dynamics predicted by RNA velocity (*see Methods*) based on exonic and intronic reads (proxies for respectively mature and nascent RNA).

To establish if other *in vivo* developmental stages were recapitulated *in vitro*, we checked the expression of SHF and FHF markers across culture methods and time points (Supplementary Fig. S7A-B). Several SHF markers were upregulated over the course of differentiation both in the atrial and ventricular lineage (*HAND2*, *ISL1* and *FOXC1*; Supplementary Fig. S7B). In the atrial EB culture conditions, an increase of *TBX5* expression occurred directly after a transient increase of *HOXA1* expression (Roux et al., 2016; Steimle & Moskowitz, 2017; Sweat et al., 2023) (Supplementary Fig. S7B). This particular expression profile is consistent with the expected posterior SHF (pSHF) identity (Protze et al., 2019). Other examples of pSHF markers that increased in the atrial EBs are *FOXF1, TBX18* and *OSR1* (Hill et al., 2019; Meilhac & Buckingham, 2018; Steimle & Moskowitz, 2017) (Supplementary Fig. S7C-D). On the other hand, the vCM cultures showed limited expression of *TBX5* in comparison to the aCM cultures (Supplementary Fig. S7B), while *FGF8* and *FGF10* increased alongside an early expression of *SIX1*, indicative of anterior SHF identity (aSHF) (Itoh et al., 2016; Yang et al., 2022) (Supplementary Fig. S7E). By examining the identity of the CMs from day 14 onwards (Supplementary Fig. S2D), we observed a higher expression of *MYL2,* together with the expression of *HAND2* and *SEMA3C* (Supplementary Fig. S7B, F) in the ventricular clusters (derived from both EB and monolayer cultures), associated with an outflow tract (OFT)-like or right ventricle CM identity (Rana et al., 2014; Yang et al., 2022). These findings suggest SHF progenitors develop in these cardiac cultures, with an aSHF identity in the ventricular culture and a posteriorizing effect of RA (pSHF) in the atrial EB culture. The ventricular cultures additionally expressed the FHF marker *HAND1* (Supplementary Fig. S7B), consistent with contributions to left ventricular CM. Collectively, these results outline multiple routes towards a cardiomyocyte fate, one with relatively low *COL3A1* expression on a mixed aSHF-FHF trajectory towards vCMs, and another of *COL3A1*-high expression progressing towards aCMs derived from a more posteriorized SHF lineage, a dynamic that is driven by treatment with RA (Supplementary Fig. S7D). Interestingly, epicardial marker genes such as *SFRP5* (Fujii et al., 2017), overlapped with *COL3A1* in the non-cardiomyocyte branches in EB cultures, irrespective of RA-induced posteriorization of the SHF (Supplementary Fig. S4C and Fig. S7G). These data document the dynamic re-routing of progenitors along developmental trajectories depending on the culture method, raising the question what molecular mechanisms are underlying this plasticity of multilineage cardiac differentiation.

### Identification of candidate cardiac regulators by integrative multiomic analysis

We focused first on the influence of RA on lineage specification in EBs, as it provides a major switch for atrial versus ventricular CM lineage specification in these cultures, in the presence of a relatively stable number of non-CM cell types (cf. Fig. 1D). To study lineage-specific transcription factor expression patterns, an EB subset of the atlas was generated to perform differential-expression testing over time on CM-committed cell clusters (Fig. 3A, Supplementary Fig. S8A-D). This analysis outlined several known CM-specification regulators, such as ventricular *IRX3* and *HEY2*, and atrial *MEIS2*, *NR2F2/1* and *HEY1*. Additionally, atrial-enriched expression was found for *LHX2*, *ZNF711* and *ZNF503* (Supplementary Fig. S8E-G), genes that are not known for their role in CM lineage commitment. Additionally, while *ZNF711* showed broad expression across both lineages, RA appeared to further induce this zinc-finger gene in the atrial lineage in a temporal manner (Supplementary Fig. S8D-E). Since RA was added to the atrial cultures for 24 hours on day 4, ‘early’ genes can be identified that are only transiently induced by RA (for example *HOXA1*), whereas ‘long-term’ genes (for example *LHX2*, *ZNF711* and *ZNF503*) are induced with delayed dynamics (Supplementary Fig. S8D, H).

**Figure 3.**
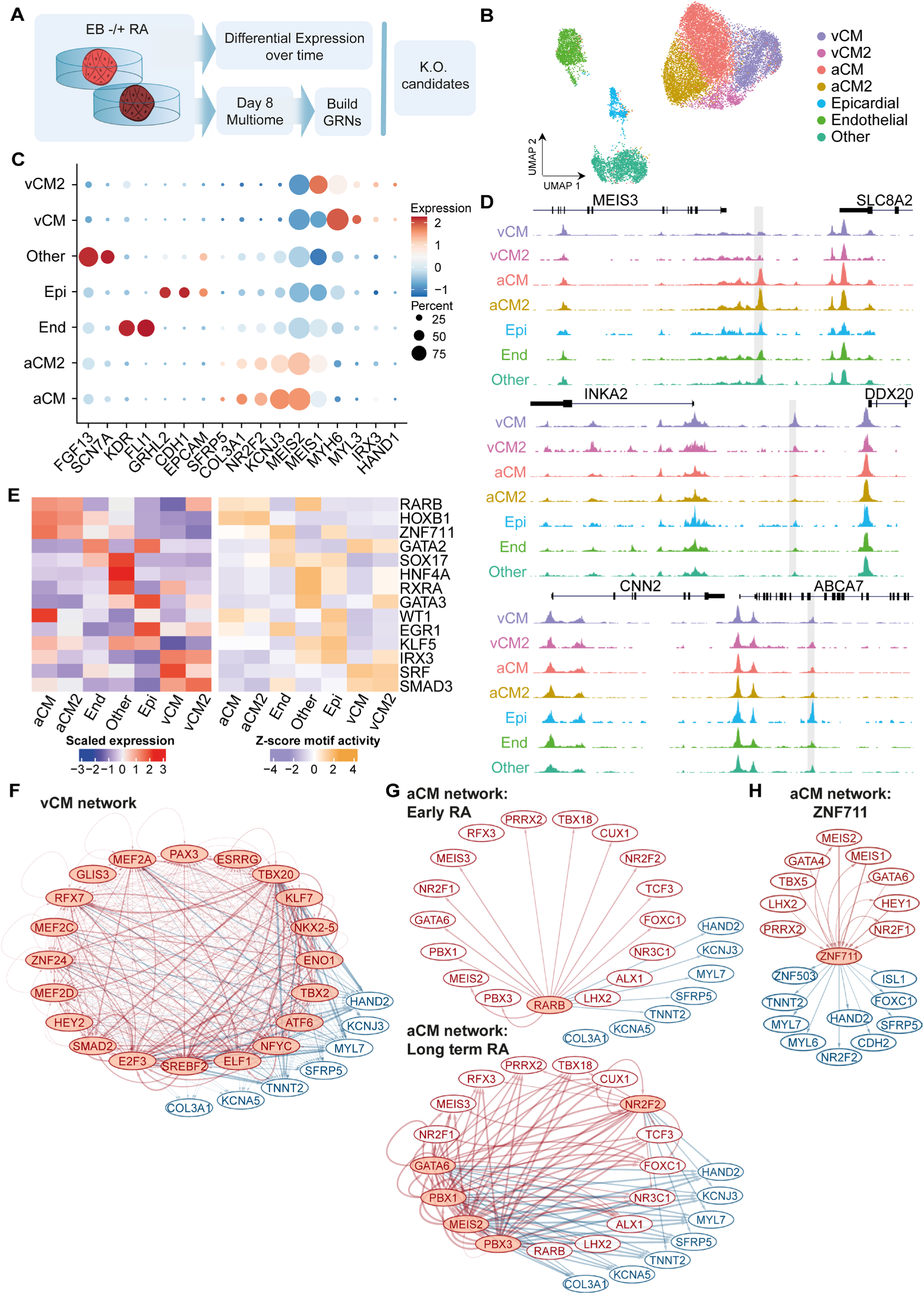
Identification of transcriptional regulators of the atrial fate with multiome data. (A) Schematic overview of the approach to select transcription factors (TFs) of interest in the +RA/non-RA-treated cultures. In brief, differentially expressed TF were determined over time (day 4 to 14) across the aCM- and vCM-committed cell clusters in the EB cultures of the roadmap, and a multiome analysis (snRNA-seq and snATAC-seq from the same nuclei) of day 8 atrial and ventricular EB cultures was used to generate cell type specific gene regulatory networks (GRNs). The combination of these approaches was used to identify potential knockout candidates. (B) UMAP representation of the multiome data from a day 8 embryoid culture (EB) directed towards the atrial and ventricular lineage. Dimensionality reduction and clustering of the cells was performed on the single-cell RNA-seq fraction of the data. (C) Scaled marker gene expression levels over the different cell states. End = endothelial; Epi = Epicardial. (D) Cluster pseudobulk signals of the scATAC-seq data, zoomed in on the three different loci in Genome Browser view, where a differentially accessible peak was identified enriched in the atrial (top), ventricular (middle) and epicardial (bottom) cluster, between the *MEIS3* and *SLC8A2* gene (top), *INKA2* and *DDX20* gene (middle) and within the *ABCA7* gene (bottom). The grey box highlights a 200bp peak identified as differential in signal across the clusters. Black boxes within the genes indicate exons. (E) Motif enrichment as identified by Maelstrom analysis on the differentially accessible peaks shown in full in Supplemental Table S1A-B and clustered in Supplemental Fig. S9D. Left heatmap shows expression levels of binding TFs, right heatmap the linked motif activity as inferred with motif analysis. (F) Differential network of the vCM over the aCM cluster, as predicted by scANANSE. Top 20 most influential transcriptional regulators (red) are shown, including 7 markers of interest (blue). (G) Differential network of the aCM over the vCM cluster, as predicted by scANANSE, again showing the top 20 most influential regulators (red) and the same targets (blue) as in (F). Nodes in the network are highlighted for the temporal expression with showing the Early RA responding gene RARB and the connections to its targets (top) and the Long-term RA responding genes and their target interactions (bottom). The highlight nodes are indicating the presence of these regulators in one of the RA gene subsets as presented in Supplemental Fig. S8H. (H) Subset of the aCM network for gene of interest ZNF711, with a selection of influential factors identified in aCM differential network (red) and targets of interest (blue).

To examine the role of these and other transcription factors at the level of chromatin accessibility, we generated an orthogonal single-nucleus multiomic dataset (snRNA-seq and snATAC-seq) of atrial and ventricular EBs at the onset of lineage commitment (day 8, Fig. 3A). Dimensionality reduction and clustering of the transcriptome modality of the data led to the identification of the cell states we identified before, with a *KCNJ3* and *NR2F1*-positive population of early aCMs, vCMs expressing *MYL7* and *MYH6,* a *GRHL2* and *CDH1*-positive population of epicardial cells, and an endothelial cluster marked by *KDR* (or *FLK1*) and *FLI1* expression (Fig. 3B-C, Supplementary Fig. S9A-C). Subclusters observed within the CM clusters (labeled vCM2 and aCM2) were marked by gene expression associated with different cell cycle phases (Supplementary Fig. S9B-C). Correlation of the multiome clusters with day 8 EB clusters of the atlas, revealed similarities of multiome clusters 4 and 8 with the other / endoderm populations of the atlas (Fig. 3B, Supplementary Fig. S10A-C), while also expressing fibroblast genes *FGF13* and *SCN7A* (Supplementary Fig. S9C).

With the clusters outlined in the multiomic data based on gene expression, peak calling and differential peak accessibility analysis was performed on the chromatin accessibility modality of the dataset. At this early stage of lineage commitment and RA-dependent change, quantitative differences in accessibility between cell clusters were observed for relatively few genes (Fig. 3D, Supplementary Fig. S9D). We then examined the transcription factor motifs correlating with differences in chromatin accessibility between cell clusters (‘motif activity’, *see Methods*), which identified motifs linked to important cardiac regulators (Fig. 3E, Supplementary Table S1A-B). For example, the motifs of RA-receptor (RARB) and HOXB1 are associated with chromatin accessibility in atrial CMs (aCM and aCM2 clusters), proteins known to be induced by RA and to be implicated in pSHF development (Roux et al., 2016). The motifs for IRX3 and SRF, important for aSHF and CM differentiation (Gaborit et al., 2012; Guo & Pu, 2020; Kim et al., 2016; Yang et al., 2022), were associated with the vCM clusters. Other examples include the WT1 and ERG1 motifs enriched in the epicardial cluster (Marques et al., 2022; Meier et al., 2023; von Gise et al., 2011), the GATA2 motif in endothelial cells (Kanki et al., 2011), and motifs for HNF4A and SOX17 in the endoderm cluster (Hanawa et al., 2017; Kanai-Azuma et al., 2002; Séguin et al., 2008). A motif for ZNF711 was associated with increased accessibility in the endothelial, epicardial and aCM clusters (Fig. 3E).

### Gene regulatory network acting downstream of retinoic acid points to LHX2 and ZNF711 as novel regulators of interest

To understand the transcriptional regulation driving CM specification in these stages of cardiac development, we predicted the CM-specific gene regulatory networks (GRNs) utilizing scANANSE, a recently developed tool for identification and prioritization of key transcription factors involved in cell fate determination (Smits et al., 2023; Xu et al., 2021). In short, scANANSE prioritizes transcription factors by integration of chromatin accessibility (sequence motif content) and transcriptome data, inferring a differential network between two biological states. The differential network allows scANANSE to prioritize factors for their inferred ability to “reprogram” cells from one biological state to another. To generate these networks, we used the complete peak set as input for scANANSE, which consisted of 150,597 peaks in total (Supplementary Table S3A). The predicted aCM network (Supplementary Table S1C), highlighted many regulators known for their role in cardiac development and regulation of cardiac progenitors from the pSHF, like *TBX18*, *MEIS2* and *NR2F2* (Lescroart & Zaffran, 2018; Quaranta et al., 2018). The vCM network included multiple known CM-regulators, such as *TBX20*, *MEF2A/C* and *NKX2-5*, but also *HEY2*, a regulator particular to the ventricular subtype (Buckingham, 2015; Churko et al., 2018) (Fig. 3F, Supplementary Table S1D). For the non-CM clusters which consisted of cells from both culture conditions, we compared the cell type specific network to an average network of all cells together (*see Methods*). Top influential factors identified this way, for the epicardial network included *GRHL2*, *TFAP2A*, *BACH2*, for endothelial network these included *ERG*, *ETS1*, *FLI1*, *ELK3*, and for the endoderm network *RFX6*, *GATA3*, *HNF4A* and *HNF4G* (Supplementary Table S1E-G).

Subsequently, we used the temporal gene expression information from our transcriptome atlas (Supplementary Fig. S8H), to infer a temporal regulatory hierarchy within the aCM network. Based on these expression dynamics within the top-ranked network of transcription factors, RA receptor beta (RARB) emerged as the regulatory starting point (Fig. 3G, top), consistent with the use of RA to trigger atrial lineage commitment. RARB is predicted to regulate multiple well-known pSHF genes (i.e. *MEIS2*, *TBX18*, *NR2F2* and *FOXC1*). Top-regulators encoded among the long-term RA-induced genes are *GATA6*, *MEIS2*, *PBX3*, *PBX1* and *NR2F2* (Fig. 3G, bottom). The predicted regulatory interactions of seven genes of interest (*COL3A1, KCNA5, TNNT2, SFRP5, MYL7, KCNJ3, HAND2*) suggest that most targets can be regulated by RARB directly, whereas *COL3A1* and *KCNA5* may be regulated further downstream in the network by the long-term regulators.

Among the influential factors of the aCM network, also ZNF711 and LHX2 were identified. Multiple influential factors were predicted to regulate *ZNF711* in the aCM network (Fig. 3H), among which known pSHF factors TBX5, HEY1, MEIS1/2 and NR2F1, but also factors more generally implicated in cardiac development such as GATA4/6 (Xin et al., 2006). ZNF711 was predicted to regulate some of these and other SHF regulators like *ISL1* and *FOXC1* and aCM-specific markers (*NR2F2*, *MYL6)*. Interestingly, ZNF711 was also predicted to regulate general CM genes, i.e. *TNNT2*, *MYL7* and *N-cadherin* (*CDH2*). Additionally, *ZNF503* was found downstream of ZNF711 in the aCM network. These findings were especially interesting since limited information is available regarding the potential role of these zinc-finger proteins in cardiac development. The highly conserved *ZNF503* protein has been shown to transcriptionally repress *GATA3* (Shahi et al., 2017) and to play a critical role during embryonic development in mice and zebrafish (Boobalan et al., 2022; Nakamura et al., 2008; Shahi et al., 2017). Our network predictions and the temporal and RA-induced expression patterns of *ZNF711* and *ZNF503* prompted us to test the function of these little known zinc finger factors in cardiac differentiation.

### Ablation of ZNF711 halts cardiomyocyte differentiation and favors the epicardial and endodermal lineages

We selected three genes (*LHX2*, *ZNF711* and *ZNF503*) to assess their roles in cardiac lineage commitment. For each of these factors a CRISPR knockout (KO) was performed in the dual NKX2.5^EGFP/+^COUP-TFII (NR2F2)^mCherry/+^ hPSC reporter line (Fig. 4A), with KO efficiencies over 70% each, both before and after cardiac EB differentiation (Supplementary Fig. S11A-B). KO hPSCs were immediately differentiated in EB cultures over the course of 14 days towards the ventricular (non-RA) and atrial (+RA) cardiac cell types. The efficiency of CM differentiation was assessed using NKX2-5-GFP reporter expression (Fig. 4B). In the ventricular *ZNF711*-KO culture, reporter expression was significantly lower compared to control (*TRAC*), while this was not observed in the RA-treated (atrial) *ZNF711*-KO EBs or in any of the other conditions. The lower CM differentiation efficiency in the ventricular *ZNF711*-KO was accompanied by an increase in the abundance of NKX2-5-GFP-negative and NR2F2-mCherry-positive fraction compared to control (Supplementary Fig. S11C-D), suggesting a shift in lineage commitment.

**Figure 4.**
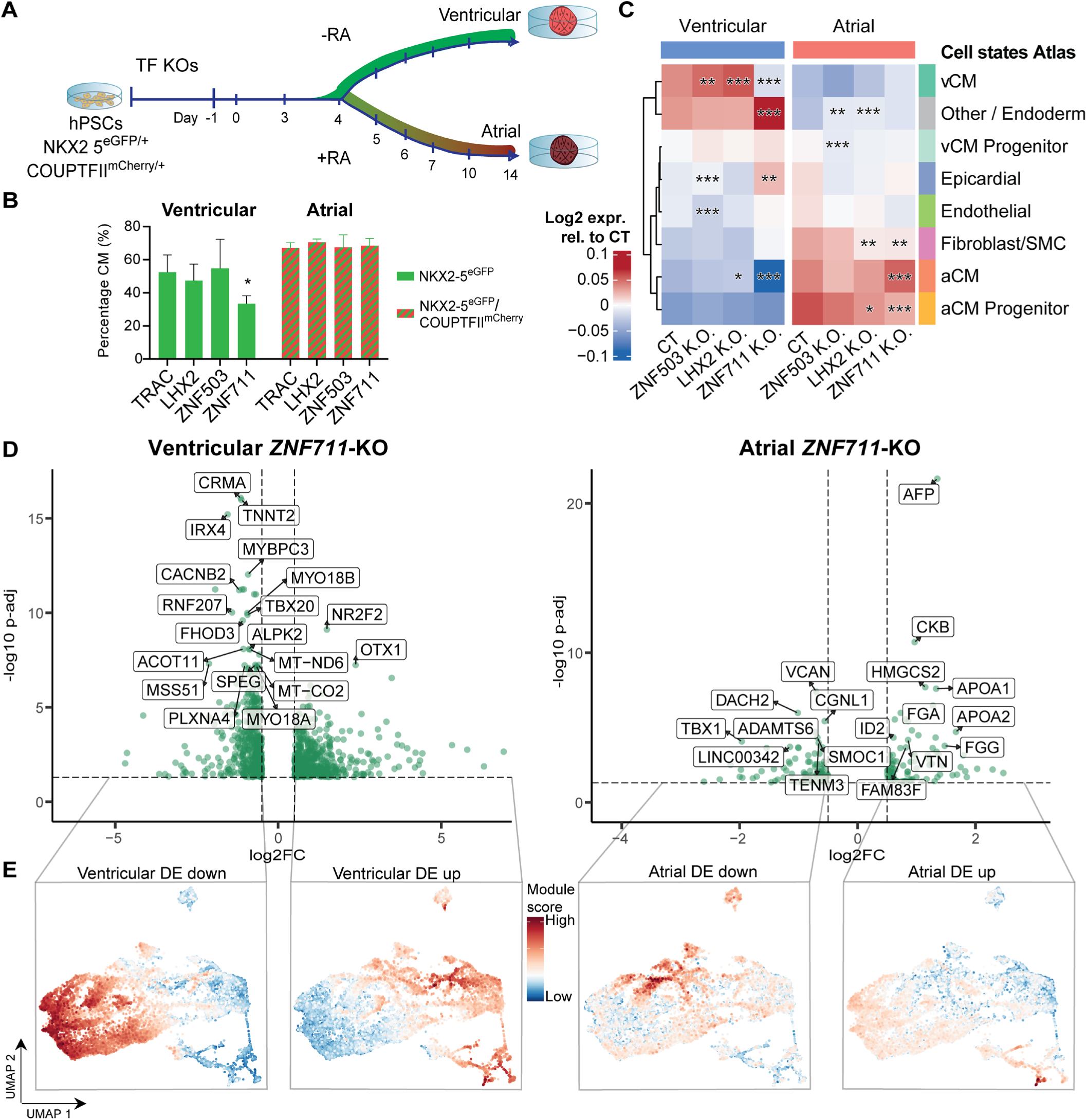
Knock-out experiments for LHX2, ZNF711 and ZNF503. (A) Experimental set-up for the transcription factor (TF) knockout (KO) experiments. hPSC carrying the *TRAC, ZNF711*, *ZNF503* or *LHX2* knockout (TF KOs) were differentiated towards aCM- and vCM-EBs (+/- RA, respectively). At day 14 of differentiation, EBs were collected for subsequent analysis. (B) Percentages of cardiomyocytes (CM) per KO, for non-RA-treated (ventricular, left green bars) and +RA-treated (atrial, green and red bars) conditions, measured in cells positive for the CM reporters NKX2-5 in non-RA-treated and NR2F2 (aCM) in +RA-treated cultures at day 14. *TRAC-*KO is the gene CRISPR control. Data are mean ± s.e.m; ordinary two-way ANOVA with Tukey’s multiple comparisons test; **p-adjusted < 0.001, *vs TRAC*. (n=3 independent differentiations per culture protocol and KO target). (C) Averaged expression scores within the *LHX2*-KO and *ZNF711-*KO compared to control. The top 200 marker genes per cell type were determined in day 14 embryoid body cells from non-RA and +RA conditions (Supplementary Table S2A), and used to average the expression levels in the CT and KO samples. Averaged expression scores are divided by the average score of both CT for the same cell type, to enable direct comparison between all conditions shown. Significance was calculated over the averaged expression levels across 200 genes, and per matched replicate and control, stars indicate the significance of the highest p-value found amongst the replicates (Methods). (*p-adjusted < 0.05, **p-adjusted < 0.001 and ***p-adjusted < 0.0001). (D) Volcano plot of the differentially expressed (DE) genes of the *ZNF711*-KO cells cultured towards ventricular cardiomyocytes (non-RA, left) and atrial cardiomyocytes (+RA, right) (adjusted p-value < 0.05). Top hits with the lowest adjusted p-value are indicated with text labels. The vertical dotted line indicates a threshold of |log2FC| > 0.5 and the horizontal dotted line |-log10P-value| < 0.05. (E) The *in vitro* atlas (Fig. 1) labelled with compound gene expression levels (module scores) based on the gene groups from left to right: non-RA downregulated (log2FC < 0), non-RA downregulated (log2FC > 0), +RA downregulated and +RA upregulated genes, based on significance (p-adjusted < 0.05) after *ZNF711*-KO. CT = control, aCM = atrial, vCM = ventricular, RA = retinoic acid, DE = differentially expressed, log2FC = log2 (expression fold change between KO and CT).

A substantial differentiation defect in the ventricular culture was corroborated by transcriptome analysis, which we performed for both the atrial and ventricular cardiac EB cultures of each KO at day 14 of differentiation. The strongest changes in gene expression were detected in the *ZNF711-* and *LHX2*-ablated cultures relative to controls (Supplementary Fig. S11E). To explore changes in cell type commitment, sets of 200 marker genes per cell type were defined from the atlas (Supplementary Table S2A) to calculate average cell type-specific marker expression levels per KO and culture condition (Fig. 4C). This analysis uncovered that changes in cell lineage-specific markers were highly dependent on the treatment with RA. The *ZNF711*-KO samples showed a significant decrease in both aCM and vCM markers under ventricular conditions (Wilcoxon p-adjusted 2.7 x 10^-9^ and 2.2 x 10^-11^, respectively). In RA-treated atrial cultures, however, vCM marker expression was not affected significantly, while the expression of aCM markers increased significantly (Wilcoxon p-adjusted 7.8 x 10^-5^) and the expression of aCM progenitor markers was reduced (Wilcoxon p-adjusted 2.6 x 10^-11^). In ventricular cultures, definitive endoderm marker expression was increased (Wilcoxon p-adjusted 1 x 10^-8^), alongside epicardial gene expression (Wilcoxon p-adjusted 4.6 x 10^-3^). The *ZNF711-*KO samples exhibited significant downregulation of general cardiomyocyte markers (*TNNT2*: p-adjusted 1.1 x 10^-16^, *MYL7*: 1 x 10^-6^, *CDH2*: *0.038, IRX4*: 6.2 x 10^-16^, *TBX20*: 1.3 x 10^-10^, *PLXNA4*: 6 x 10^-8^) (Fig. 4D,

Supplementary Table S2B) and gene ontology terms (Supplementary Table S2H) in the ventricular cultures. This was not observed in the RA-treated atrial *ZNF711*-KO samples (Supplementary Table S2C), in which upregulated genes were enriched for the terms *muscle cell differentiation* and *muscle system process* (Supplementary Table S2H). This suggests that the inhibition of cardiomyocyte differentiation observed in ventricular cultures, was rescued by RA in the atrial *ZNF711*-KO cultures. To characterize this in more detail, *ZNF711-*KO up- and down-regulated genes were used to calculate module scores in the single-cell *in vitro* atlas (Fig. 4E). The ventricular *ZNF711-*KO downregulated targets (Fig. 4E, left panel) normally show high expression in all of the CM-committed clusters within the atlas, whereas the ventricular *ZNF711*-KO upregulated genes are normally expressed in the non-CM branches of the atlas (Fig. 4E, middle-left). These results were corroborated by the increased NR2F2 protein levels measured in a NKX2-5-negative population (Supplementary Fig. S11C-D), combined with increased expression levels of *COL3A1* and *SFRP5* (p-adjusted 0.029 and 0.019, respectively) (Supplementary Table S2B) in these ventricular *ZNF711*-KO cultures compared to control, indicating a shift in populations from CMs to non-CMs. Conversely, the atrial *ZNF711*-KO downregulated genes were mostly expressed in the non-CM populations and CM progenitors (Fig. 4E, middle-right) and upregulated genes showed specific expression in later CM differentiation stages of the atlas (Fig. 4E, right panel). These data show that *ZNF711*-KO hampered CM differentiation and increased both progenitor and non-CM signatures in the absence of RA, whereas this effect was completely rescued by administration of RA.

Differential expression testing in the *ZNF503*-KO EBs resulted in a more limited number of differentially expressed genes in either condition (Supplementary Table S2D-E). *LHX2*-KO showed an increase in vCM and aCM markers (Wilcoxon p-adjusted 1.8 x 10^-2^ and 1.2 x 10^-8^, respectively) in the ventricular EB cultures (Fig. 4C, Supplementary Table S2F-G). In the atrial cultures, a significant decrease in fibroblast / smooth muscle markers (Wilcoxon p-adjusted 3.1 x 10^-3^) and aCM progenitor markers (Wilcoxon p-adjusted 3.5 x 10^-2^) was observed, to the benefit of the ‘other’ cluster (Wilcoxon p-adjusted 3.4 x 10^-4^), marking a substantial defect in differentiation in combination with RA treatment.

### Targets under combined regulation of RARB and ZNF711 confirm predicted regulatory network

Given this dynamic re-routing of cardiac progenitors along different lineages with these genetic perturbations, we wondered how the observed transcriptomic changes related to both the gene-regulatory networks and the transcription factor motif content in our chromatin accessibility data. We therefore determined the overlap between *ZNF711*-KO differentially expressed genes (DE-genes) and the ZNF711 targets predicted by scANANSE (Fig. 5A and Supplementary Fig. S12A, Supplementary Table S3B, *see Methods*). Since scANANSE generates cell type-specific GRNs, we performed this test for the vCM, aCM and epicardial networks. The scANANSE ZNF711 targets were enriched for the ventricular *ZNF711-*KO DE-genes, in both the vCM and aCM networks (fold enrichment 1.4 and 1.9, hypergeometric p-adjusted 5 x 10^-05^ and 2.4 x 10^-12^ respectively), but not in the epicardial network, whereas this enrichment was not found for the overlap with the DE-genes in the RA-treated (atrial) condition in any of the cell networks (Fig. 5A). These findings confirm the predictive power of our scANANSE inferred networks. This also suggests that *ZNF711*-KO is directly required for the shared atrial-ventricular CM program, but that RA rescues this program while also conferring an atrial identity to the resulting CMs.

**Figure 5.**
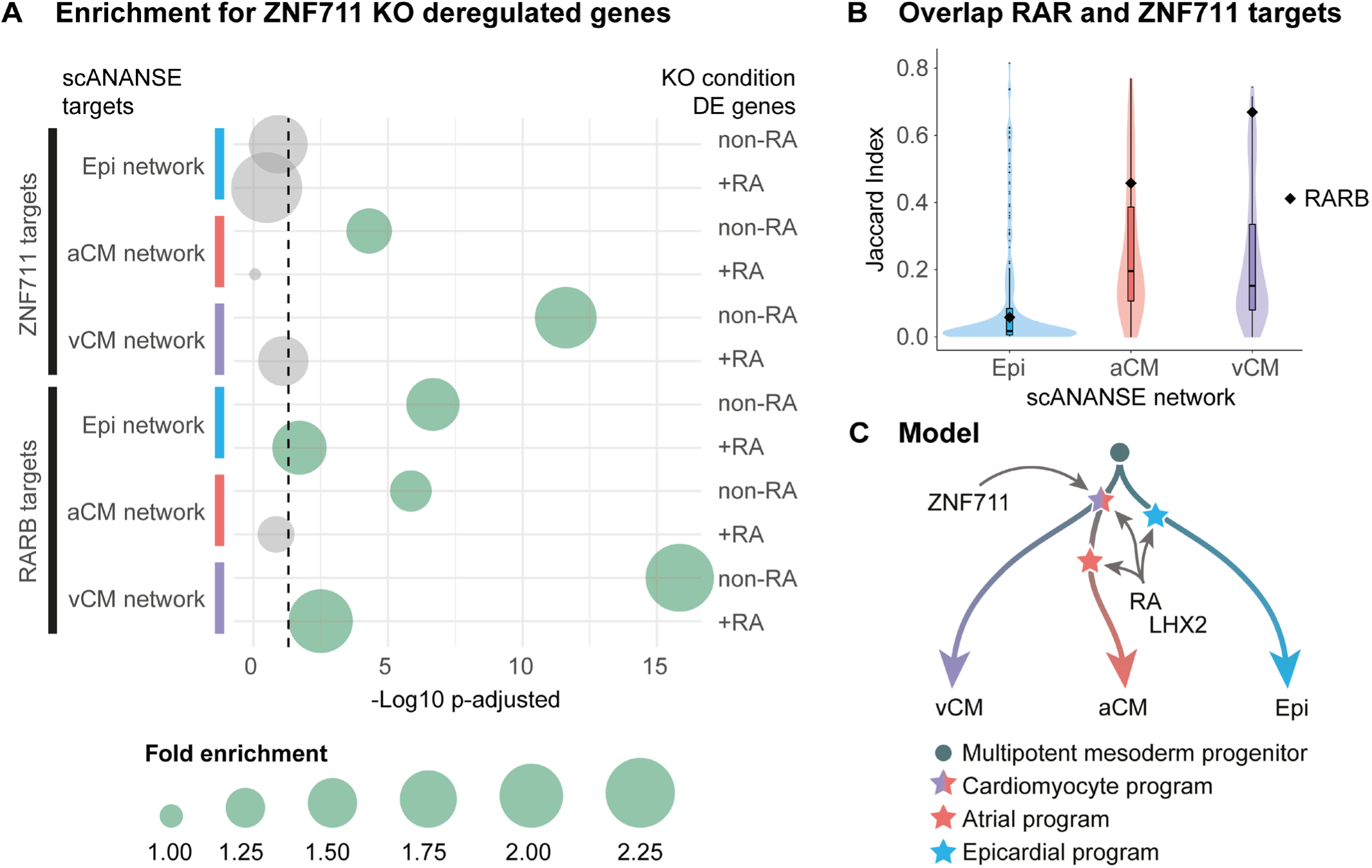
Enrichment for ZNF711 targets in the scANANSE networks and proposed model. (A) Enrichment of ANANSE-predicted network targets among differentially expressed genes in ZNF711 knockout cardiac embryoid bodies. Bubble size represents fold enrichment, whereas the X-axis shows multiple-testing corrected hypergeometric test p-values. Indicated on the left of the bubble plot is the regulator for which the predicted targets were selected from each of the cell type-specific scANANSE networks. On the right of the plot, the conditions are indicated from which the *ZNF711*-KO differential genes were selected, non-RA (ventricular) or +RA (atrial). The dashed line visualizes the p-adjusted value of 0.05. This figure represents a subset of all comparisons. See Supplementary Figure S12 and Supplementary Table S3B for all comparisons, including results on LHX2. (B) Violin plots showing the overlap (Jaccard Index) between cell type-specific ANANSE-predicted targets of ZNF711 and RARB and other transcription factors (violin plots). The black diamonds indicate the fold enrichment of overlap between ZNF711 and RARB targets for each of the networks. See Supplementary Table S3C-E for the table with results. (C) Model of requirements for balanced lineage commitment of progenitors to epicardial cells, atrial and ventricular cardiomyocytes by the concerted action of *ZNF711*, *LHX2* and retinoic acid.

Interestingly, performing the same test with the targets predicted by scANANSE for RARB, showed significant enrichment of RARB targets in all cell types among the DE genes in the ventricular (non-RA) *ZNF711*-KO cultures (Fig 5A and Supplementary Fig. S12A, Supplementary Table S3B). To establish how this overlap between RARB and ZNF711 targets compares to other regulators, we determined the overlap by pair-wise calculation of the Jaccard Index (number of shared targets divided by the number of the union of targets of two transcription factors) of ZNF711 predicted targets with the targets of other transcription factors, and compared the distribution of Jaccard Indices with the index of the ZNF711-RARB pair (Fig. 5B). In each case, just over 200 transcription factors constitute the cell specific network, each predicted to regulate up to ∼19,000 target genes, with a probability > 0.8 (Supplemental Table S3C-E). These works are highly interconnected (as visualized for the top 20 transcription factors, cf. Fig. 3F-G), and many show extensive overlap in targets (Fig. 5B). ZNF711 showed a large overlap with RARB targets compared to other regulators in this network, especially within the vCM network (Jaccard Index 0.67, 26-fold-enrichment, hypergeometric p-adj. < 1e-100). In the aCM network these shared were also enriched, although to a lesser degree (Jaccard Index 0.46, 8.8-fold, hypergeometric p-adj. < 1e-100). These shared targets can explain how RA-treatment can rescue CM differentiation in *ZNF711*-KO atrial EB cultures.

These analyses also showed significantly enriched overlaps of LHX2 targets in the epicardial and aCM networks and *LHX2*-KO DE genes, and moreover of LHX2 targets and *ZNF711*-KO DE genes and vice versa, but only under RA-treatment conditions (Supplementary Fig. S12A-B, Supplementary Table S3B). No major shift is observed in the expression of cell type markers in RA-treated *LHX2*-KO EBs, whereas in the absence of RA, ventricular CM gene expression is stronger, and no significant effect is observed for epicardial gene expression (Fig. 4C). This suggests that LHX2 cooperates with RA receptors to activate epicardial and CM gene expression.

As summarized in our model (Fig. 5C), the data uncover how the balance of lineage decisions of cardiac progenitors is shifted by genetic perturbations, and is modified in major ways by RA, a natural cardiac crescent morphogen. We find that *ZNF711* is important for a general CM program including the expression of structural sarcomere genes, but only in the absence of RA. The resulting block in CM differentiation redirects progenitors towards the epicardial and endodermal lineages. RA rescues the defect on the general CM program in the absence of ZNF711 and induces the atrial CM identity.

## DISCUSSION

We present a roadmap of *in vitro* cardiac differentiation, detailing the steps of lineage commitment and differentiation in cardiac monolayer, EB and EHT cultures. This allows for direct comparisons over time and between anterior or posteriorized SHF progenitors. We identified similar multilineage compositions within the atrial and ventricular EB cultures as reported in other cardiac organoid protocols (Meier et al., 2023; Silva et al., 2021). In our experiments, the EBs showcased a plasticity of multipotent progenitor cells that was susceptible to patterning by RA. Because of this high level of plasticity in cardiac cultures and the ability to culture CMs of various specific subtypes (Devalla et al., 2015; Schmidt et al., 2023; Schwach et al., 2017; Zawada et al., 2023), detailed deconstruction of the developmental steps within the culture is necessary, to understand and optimize the cell type composition. The specification of posterior SHF progenitors directed CMs towards an atrial identity, as expected; however, other cell lineages (endothelial, epicardial, fibroblast and SMC-like cells) were minimally affected. Our analysis of the *in vitro* roadmap not only identified known and candidate regulators, but also provides insights when tracing back and interpreting results from knockout studies. Such analysis revealed that *ZNF711* plays a critical role in safeguarding the general CM program, whereas in the absence of both *ZNF711* and RA a major shift towards the epicardial and endodermal lineage is observed.

Epicardial and anterior endoderm cell populations have been observed in previous studies (Drakhlis et al., 2021; Hofbauer et al., 2020; Meier et al., 2023; Silva et al., 2021; Zawada et al., 2023). Our findings corroborate these observations and demonstrate that these populations can emerge independently of RA treatment (Drakhlis et al., 2021; Silva et al., 2021). This was confirmed by the expression of markers of the juxta-cardiac field and pre-epicardial cells (Meier et al., 2023), including the expression of the newly identified epicardial markers such as *ITGA8* (Zawada et al., 2023). Paracrine signaling from endodermal and epicardial populations has been reported to be beneficial for both the epicardial and CM cell populations (Quijada et al., 2020). Moreover, anterior endoderm is essential for vertebrate heart development and morphogenesis (Nascone & Mercola, 1995; Withington et al., 2001). As we further characterized our multipotent progenitors around day 8, we identified potential transcriptional regulators that could steer these dynamic cultures to the specific cardiac subtypes. This led to cell type-specific GRNs that represent both known and hitherto unknown regulators of the cardiac development.

In our pursuit of candidate regulators, we experimentally identified an important role for *ZNF711* a gene about which comparatively little is known. Mutations in *ZNF711* have been associated with X-linked intellectual disability and an altered DNA methylation signature in blood (Wang et al., 2022). *ZNF711* is a member of a small family of zinc-finger genes that encode a similar N-terminal acidic activation domain (Mardon et al., 1990; Ni et al., 2020). Moreover, ZNF711 can bind CpG island promoters where it can recruit PHF8, a histone H3K9 demethylase (Kleine-Kohlbrecher et al., 2010). Notably, *ZNF711* orthologs are found in the genomes of jawed vertebrates, but not in those of invertebrate animals. Although shared among most bilaterian animals, the heart has evolved significantly in vertebrates, including the addition of chambers and the endocardial and epicardial layers (Carmona et al., 2010; Simões-Costa et al., 2005). The RA pathway was co-opted to derive atrial cardiomyocytes and the epicardial lineage from mesodermal precursors. This may have required changes in the CM specification program to safeguard the CM program. Our research reveals that *ZNF711* is important for cardiomyocyte differentiation in the absence of RA. In scenarios where both ZNF711 and RA are absent, there is a shift towards the differentiation of epicardial cells at the expense of ventricular cardiomyocytes. Cardiomyocyte commitment is rescued by RA, while simultaneously endowing the cardiomyocytes with a posteriorized atrial gene expression identity. Consistent with this rescue, we find that regulatory elements of the genes involved contain motifs for both ZNF711 and retinoic acid receptors, providing a mechanism for balanced lineage commitment both in the absence and the presence of RA. Given the plasticity of cardiac mesoderm and heart field progenitors, which are readily redirected to different cell lineages, safeguards are necessary for such a balanced lineage differentiation, and ZNF711 appears to have an important role in securing the cardiomyocyte program.

The cardiac organoid models and CM differentiation protocols available to date, have been obtained by extensive experimentation and diligent optimization. These models contain different cell types and cell states, reflecting both lineage potential and culture conditions. Further effort will be required to improve these human stem cell-based models by leveraging single-cell roadmaps and gene-regulatory networks generated by us and others. This will increase our fundamental understanding of the heart as a complex tissue, while providing a major impetus for analyses of heart disease mechanisms, heart regeneration and drug screening.

## METHODS

### HPSC culture

Experiments were performed using the human embryonic stem cell (hESC) lines NKX2.5^EGFP/+^COUP-TFII^mCherry/+^ (Schwach et al., 2017). hPSCs were maintained as undifferentiated colonies in Essential 8 medium (Thermo Fisher, A1517001) on vitronectin (Thermo Fisher, A31804)-coated 6-well plates (Greiner Bio-One, 657160).

### Differentiation of hPSCs into ventricular monolayers

Differentiation of hPSCs to VM monolayers was performed as previously described (Birket et al., 2015). Briefly, one day before starting the differentiation, hPSCs were seeded at a density of 20-25×10^3^ cells per cm^2^ on Matrigel (83 μg protein/mL) (Corning, 354230)-coated 6-well plates in Essential 8 medium. After 24 h (day 0), mesodermal differentiation was induced by addition of Activin-A (20 - 30 ng/mL, Miltenyi 130–115-010), BMP4 (20 - 30 ng/mL, R&D systems 314-BP/CF) and WNT activator CHIR99021 (1.5 - 2.25 μmol/L, Axon Medchem 1386) in BPEL medium (Ng et al., 2008). At day 3, BPEL containing WNT inhibitor XAV939 (5 μmol/L, R&D Systems 3748) was used to refresh cells. On day 7, 10 and 14 cells were refreshed with plain BPEL.

### Differentiation of hPSCs into ventricular and atrial EBs

Differentiation of hPSCs to ventricular or atrial EBs was performed as previously described (Schwach et al., 2022). Briefly, hPSC were seeded and aggregated in EB formation medium (Essential 8 containing 400 µg/mL Poly (vinil alcohol) (Sigma Aldrich, P8136) and 10 µM Y-27632 (Bio-Connect, S1049)) one day before starting the differentiation. After 24 h (day 0), mesodermal differentiation was induced by addition of Activin-A (30 ng/mL, Miltenyi 130–115-010), BMP4 (30 ng/mL, R&D systems 314-BP/CF), WNT activator CHIR99021 (1.5 μmol/L, Axon Medchem 1386), VEGF (30 ng/µL, Miltenyi Biotec, 130-109-386) and SCF (40 ng/µL, Stem Cell Technologies, 78062.1) in BPEL medium (Ng et al., 2008). On day 3 of differentiation, EBs were refreshed with plain BPEL. On day 4, atrial EBs were refreshed with either 1 µM of all trans RA (Sigma Aldrich, R2625) or 20 µM RARα selective agonist BMS-753 (Torcis Bioscience, 3505). Next day, atrial EBs treated with BMS-753 were refreshed with plain BPEL. EBs at day 7 were plated on 0.1% gelatin coated wells in BPEL medium. EBs were refreshed again with BPEL at day 10. Unless indicated, EBs at day 14 were switched to cardiomyocyte media containing 3ʹ,5-Triiodo-L-thyronine sodium salt (T3), Dexamethasone (D) and Long R3 IGF-I human (IGF, I) (CM-TDI) (Birket et al., 2015) and refreshed every 2-3 days until used at day 21.

### Generation of engineered heart tissues

Ventricular monolayer, ventricular EBs and atrial EBs were sorted at day 14 to separate CMs positive cells (NKX2.5^EGFP/+^-positive COUP-TFII^mCherry/+^-negative for vCMs and NKX2.5^EGFP/+^-positive COUP-TFII^mCherry/+^-positive for aCMs) and non-CMs (NKX2.5^EGFP/+^-negative). For each CM differentiation (ventricular mono and EBs, and atrial EBs), cells were mixed back at a specific ratio of 80% CMs + 20% non-CMs. EHTs from ventricular monolayers and atrial EBs were generated by following the protocol as described (Ribeiro et al., 2022). As a 2D control, monolayers from each CM differentiation were generated by seeding the cells with the same cell ratio (80% CMs + 20% non-CMs) in vitronectin coated 48 wells plates. EHTs and 2D controls were kept in CM-TDI media and refreshed every 2-3 days until use.

### Immunohistochemistry and confocal imaging

For whole mount staining on intact EBs, EBs at day 7 were seeded on gelatine-coated glass bottom plates. Cells were fixed at day 21 with 4% Paraformaldehyde for 15 min at RT. Permeabilization was followed with 0.1% Triton X100 for 8 min at RT. After blocking for 1 h at RT with 5% Fetal Bovine Serum (FBS), cells were incubated at 4°C overnight with the primary antibodies (Table 1). After washing, cells were incubated for 1 h at RT with secondary antibodies (Table 1) and 5 min at RT with DAPI. EHTs at day 12 after tissue formation were fixed with 4% Paraformaldehyde for 1 h at RT. Whole tissue immunostaining was performed as described (Ribeiro et al., 2022). Antibodies used are described in Table 1. Confocal images were captured using a Zeiss LSM 880 microscope.

**Table 1.** Antibodies used for immunohistochemistry on EBs and EHTs.

| Antibody | Cat. Number | Dilution factor |
| --- | --- | --- |
| Rabbit anti-NKX2.5 | Cell Signaling, 8792 | 1:400 |
| Mouse anti-COUP-TFII | Perseus Proteomics, 1:500 | 1:200 |
| Mouse anti-ACTN2 | Sigma, A7811 | 1:800 |
| Rabbit anti-MYL7 | R&D Systems, MAB90961-SP | 1:100 |
| Rabbit anti-MYL2 | Sigma HPA019763 | 1:100 |
| Mouse anti-cTnT | ThermoFisher MA512960 | 1:400 |
| DAPI | Thermo Scientific, D1306 | 1:2000 |
| Chicken anti Rabbit IgG (H+L) Alexa Fluor 488 | Invitrogen, A-21441 | 1:500 |
| Donkey anti Mouse IgG (H+L) Alexa Fluor 647 | Invitrogen, A-31571 | 1:500 |

### Optical membrane potential imaging

For optical membrane potential imaging, aCMs or vCMs at day 14 were stained for 30 min at RT with the voltage-sensitive dye FluoVolt™ according to manufacturer. For signal acquisition, the loaded sample was excited using a 488 nm laser and recorded for 10 seconds. Movies were acquired with a Nikon ECLIPSE Ti2 fluorescent microscope under temperature and humidity control (37°C and 5% CO_2_). Videos were analyzed with BV Workbench, Brainvision Inc.

### FACS and flow cytometry

Monolayers and EBs from day 3 to day 8 were dissociated to single cells with TypLE 1X (Thermo Fisher, 12563029) for 5 minutes at 37°C. From day 8 onwards, monolayers and EBs were dissociated with TypLE 10X (Thermo Fisher, A1217702) for 10 min at 37°C. EHTs were dissociated with collagenase type II (240U/ml) (Worthington Biochemical LS004176) for 20-30 min at 37°C followed by TrypLE 10X for 10-15 min at 37°C. Cells were transferred to FACS buffer (2mM EDTA, 0.5% BSA in DPBS) and filtered (VWR, 734-0001). For single cell RNA sequencing, cells were single cell sorted into 384-well primer plates. Sorting was performed with a Sony SH800. Flow cytometry was performed with a MACSQuant Analyzer 16 (Miltenyi). Flow cytometry data was analyzed with FlowJo software (Miltenyi).

### CEL-Seq2 RNA-sequencing sample preparation

Libraries were prepared in 384-well plate format using a modified SORT-Seq, or CEL-Seq2, protocol (Hashimshony et al., 2016; Muraro et al., 2016). Primer plates were generated with CEL-Seq2 compatible oligo’s containing the T7 promoter, a 5’ Illumina adapter, unique molecular identifiers (UMIs; 8 bp), a unique cell barcode (8 bp) and oligo(dT)N for mRNA molecules tagging. The primers were designed reversed from classic CELSeq2 primers, leading to barcode and UMI priming in read 2, where the transcripts were sequenced from read 1, as previously described (Gerlach et al., 2019). After sorting, the plates were spun down and immediately stored at −80°C and thawed before the reverse transcription PCR (RT-PCR). ERCC RNA Spike-In Mix II (Invitrogen) was added in 1:50 x 10^3^ ratio to each of the wells administered in 150 nL volume by the micro-dispenser Nanodrop II (BioNex). The Nanodrop II was additionally used for dispending the reverse transcription mix, consisting of Superscript II (Invitrogen) and RNasein Plus (Promega), as well as for the second strand synthesis with a mix of E. coli RNase H (Invitrogen), E. coli DNA polymerase and E. coli DNA ligase (New England Biolabs). After the second strand synthesis the cDNA of all single cells of a plate were pooled, before performing an overnight in vitro transcription reaction at 16°C using the MegaScript IVT kit (Invitrogen). Remaining primers were removed by using EXOI/rSAP-IT (Applied Biosystems) and the amplified RNA was chemically fragmented and cleaned up with AMPure XP beads (Beckman Coulter). Using random octamers the following RT was performed, after which the library amplification was one with Phusion High-Fidelity DNA Polymerase (New England Biolabs) during which the Illumina adapters were introduced, as well as the unique index sequence per library (NextFlex DNA barcodes, PerkinElmer). After bead-purification, the library concentration and quality were assessed using the DeNovix and BioAnalyzer High Sensitivity DNA Kit (Agilent Technologies), respectively. Sequencing was performed with the NextSeq 500 (Illumina).

### 10X Multiome sample preparation

Nuclei from atrial and ventricular EBs at day 8 of differentiation were isolated with an ice-cold lysis buffer consisting of 10 mM Tris-HCl (pH 7.4), 10 mM NaCl, 3 mM MgCl2, 1% BSA, 0.1% Tween-20, 0.1% Nonidet P40 Substitute, 0.01% Digitonin, 1 mM DTT (Invitrogen, cat. no. 18064-014), 1 U/µL Protector RNase inhibitor (Roche, 3335399001) in Nuclease-free water. After three washes with ice-cold nuclease-free water containing 10 mM Tris-HCl (pH 7.4), 10 mM NaCl, 3 mM MgCl2, 1% BSA, 0.1% Tween-20, 1 mM DTT and 1 U/µL RNase inhibitor, isolated nuclei were kept on ice in PBS containing 0.04% BSA and 1 mM MgCl2. The number of nuclei was counted with an hematocytometer. Droplet formation and library generation was performed as described using the 10X Genomics Next Gem Single Cell Multiome ATAC + Gene Expression Reagents kit.

### Generation of CRISPR-Cas9 knockouts for LHX2, ZNF711 and ZNF503

hPSCs maintained in StemMACS™ PSC-Brew XF (Miltenyi Biotec, 130-127-865) were dissociated as the regular maintenance passaging procedure. Per knockout target, cell pellets of 200×10^3^ cells were resuspended with 20 µL of supplemented nucleofector solution P3 Primary Cell 4D-Nucleofector® S (Lonza, V4XP3032) with indicated amounts of multi-guide sgRNAs (Synthego) and SpCas9-2NLS Nuclease (Synthego) in a ratio of 3:1, and nucleofected with a 4D-Nucleofector Core and X unit (Lonza) using program CB-150. Human *TRAC* sgRNA was used as a positive control, as recommended by the supplier. After nucleofection, cells were seeded in 2 wells of a vitronectin-coated 24-well plate with StemMACS™ PSC-Brew XF supplemented with 1:200 RevitaCell (Thermo Fisher, A2644501). Cells were kept for 2-3 days. During passaging of each well, cells were expanded and lysed for PCR amplification and Sanger Sequencing. Two passages after nucleofection, differentiation towards atrial and ventricular lineages was initiated. After atrial and ventricular differentiation, cells were characterized and lysed for PCR amplification and Sanger sequencing. Knock-out efficiency was evaluated with the ICE CRISPR Analysis tool (Synthego).

### Bulk RNA-sequencing sample preparation

For the transcription factor knockout validation experiments, RNA was harvested at day 14 of the protocols and purified using the Nucleo Spin RNA (Macherey-Nagel) kit according to the manufacturer’s protocol. Libraries were generated with the RiboErase depletion RNA-sequencing protocol by KAPA (Roche, 8098131702) and sequenced on the Illumina Next Seq 2000. Per sample 200 ng of total RNA was used for library preparations with the KAPA RNA HyperPrep Kit with RiboErase (HMR) (Kapa Biosystems), including steps of the oligonucleotide hybridization, rRNA depletion, DNase digestion and RNA elution, were performed as instructed by the manufacturer. Fragmentation and priming were performed at 94°C for 6 min, followed by first-strand synthesis, second strand synthesis and A-tailing, which were performed according to protocol. Library amplification was performed with 10 cycles, followed by a 0.8X bead-based cleanup. Library concentration and quality were assessed using the DeNovix and BioAnalyzer High Sensitivity DNA Kit (Agilent Technologies). Sequencing was performed with the NextSeq 500 (Illumina).

### Computational methods

#### Processing and integration of CEL-Seq2 scRNA-sequencing data

The FASTQs were trimmed and aligned with seq2science v0.4.0 (van der Sande et al., 2023). In short, kb-python v0.25.0 was used for quantifying (kb count with the setting ‘--technology 1,8,16:1,0,8:0,0,0’) to map the reads against the GRCh38 genome and quantify the UMI counts per cell (Melsted et al., 2021). Fluorescence reporter genes and ERCC sequences were added to the genome before the index was generated with kb-python. The combined single-cell gene count matrix was filtered for good quality cells (UMI >= 1,000, detected genes >= 500, mitochondrial genes percentage < 50% or ERCC count < 20%), after which 12706 cells were left for downstream analysis. Scater v1.20.1 was used to perform quality control on the cells of the dataset and check confounding variables within the metadata (McCarthy et al., 2017). Since four culture experiments were combined, the object was split per experiment (Mon, EB or EHT experiment or one experiment combining all three), to individually normalize and select highly variable features (2,000 each) per experiment and finally perform integration anchor selection with Seurat (v4.0.4) (Hao et al., 2021), to lower batch effects and create one larger atlas. Dimensionality reduction was performed with PCA and the first 50 principal components were used to generate the atlas UMAP and compute the shared nearest neighbor graphs using Jaccard index. Louvain clustering was performed with a resolution of 0.9 with the implementation FindClusters in Seurat. Optimal resolution settings for clustering, was evaluated with Clustree v0.4.3 (Zappia & Oshlack, 2018). For all gene expression visualizations onto the UMAP, the dots representing cells were plotted in the order of low to high expression.

#### Atlas integration validation

A total of 33,108 anchors (cell-pairs) were found between the experiments (2 independent differentiations per culture type). Anchor pairs from the same time point and method were counted and their scores averaged, which resulted in a per time point and method pairs scoring across the experiments (with a total of 535 of combinations). The product of the mean score and frequency of the anchor (score * (frequency anchor / total number of anchors)) was visualized with the R package pheatmap v1.0.12.

#### Comparison with *in vivo* human fetal scRNA-seq data

Two available human fetal scRNA-seq datasets from Cui et al. and Asp et al. were integrated by use of Seurat (Asp et al., 2019; Cui et al., 2019). The annotation of the Asp et al. dataset as established by the authors, was used to annotate the clusters of the integrated dataset. The first 20 principal components were used for running UMAP dimensionality reduction and neighborhood graph, and the Louvain clustering was performed with a resolution of 1. The resulting dataset was used for predicting the cell types of the in vitro atlas clusters, by running the *TransferData* functionality of Seurat on the integrated assays with the in vivo dataset as the reference. The resulting prediction scores per cell of our in vitro atlas (a calculation outlined in the paper of Stuart et al., 2019) were averaged to create a prediction score per atlas cluster and visualized with pheatmap.

#### RNA velocity on early atlas subsets

Subsets of the atlas were generated by taking the time points of differentiation until day 14 (before transfer to maturation medium) and split per protocol, resulting in three early subsets: ventricular monolayer, atrial and ventricular EB cultures from day 4 to day 14. Dimensionality reduction was performed in Seurat and UMAPs were generated with the first 30 principal components and a minimum distance set to 0.75. The raw spliced, unspliced counts, atlas cluster information and UMAP embeddings from the Seurat object, were transferred to an AnnData object for further analysis with Scanpy v1.8.2 in Python (Wolf et al., 2018). The genes selected for finding the integration anchors were taken along as HVG. RNA velocity was run with scVelo v0.2.4 (Bergen et al., 2020).

#### Embryoid body differential transcription factor analysis over time

For direct aCM and vCM comparison, a subset of only embryoid body time points was selected. Differential gene expression testing was performed on selected clusters, representing the progenitor stages contributing to aCM and vCM over time. Wilcoxon Rank Sum test was used to find significant differentially expressed genes between the clusters 24 and 6, 14 and 0, 9 and 4 and 2 and 3 (atrial and ventricular clusters, respectively), expressed in at least 25% of the cells in one of the clusters in each comparison and with an adjusted p-value < 0.05. The resulting list of genes was filtered on genes considered to be transcription factors (Lambert et al., 2018). Hierarchical clustering of the factors was performed to separate specific expression patterns over the different time points across the lineages, and the results were visualized using Complex Heatmap v2.8.0 (Gu et al., 2016).

#### Retinoic acid-related gene sets

Differential expression with Wald’s ranking test was performed for each of the time points between the lineages in the EB subset. The significantly upregulated genes in the atrial (the RA-induced condition) over ventricular (non-RA), were separated into three subsets relative to the time of RA administration. This was done in the following way: 1) Early RA genes were genes considered significantly upregulated at day 5, that were not upregulated at d4 or any of the time points after day 5 in aCM over vCM, 2) the Long Term RA genes were selected as not yet upregulated at day 4, but upregulated from day 5 onward and still upregulated at the later time points (d8 and d14), and 3) the Late RA genes were selected as genes the genes upregulated at d8 and not yet upregulated at day 4 or 5 in aCM over vCM.

#### Multiome analysis processing and integration

The multiome data was mapped with 10X Genomics Cell Ranger ARC v2.0.1 on the GRCh38 genome. To select droplets containing a quality cell, the cell-calling performed with running Cell Ranger ATAC on the ATAC data alone was combined with setting additional manual thresholds (only keeping droplets with a number of RNA counts > 1,000 and < 10,000, number of ATAC counts > 5,000 and < 20,000, nucleosome signal < 1, TSS enrichment score > 4 and percentage of mitochondrial reads < 20%). The resulting dataset consisted of 17,027 cells and was analyzed using Signac v1.7.0 and Seurat v4.0.3 (Stuart et al., 2021). The EB day 8 samples, one from each lineage, were integrated using CCA on the scRNA-seq matrices and visualized using UMAP on the principal components of PCA analysis on the top 2,000 highly variable genes of the data. Clustering was also performed on the scRNA-seq matrices. Pearson correlations between the Multiome scRNA-seq clusters and the CEL-Seq2 derived clusters of d8 EB cells, were performed on subsampled (1,000 cells maximum per cluster) datasets, and the R2 values per cell-cell comparison was visualized with pheatmap.

#### Quality control and motif analysis on scATAC-seq pseudobulk of the clusters

Per-cluster pseudobulk BED-files of the scATAC-seq modality of the multiome, were generated using the SplitFragments functionality in Signac. The BED-files were scaled and converted to BedGraph-files using *genomecov* from bedtools v2.30.0 (Quinlan & Hall, 2010), sorted, and processed to BigWig-files using the UCSC *BedGraphToBigWig* function (Kent et al., 2010). Visualization of the BigWig-files was performed in the Genome Browser (Kent et al., 2002), genes without gene name annotation were removed from this view. Peak calling was performed with the MACS2 implementation of Signac, for each of the clusters as well as the pseudobulk of all scATAC-seq reads in the dataset. The subsequent peak summits were merged in case they were at a distance of < 100 bp to one another, and replaced for a summit in the middle of those locations. To generate a merged peak set, all of the summits were extended with 100 bp up- and downstream of the summit. This peak set was used for differentially accessibility testing with the *FindMarkers* functionality from Signac using logistic regression, with the total number of fragments as latent variable. The resulting set was visualized in a bandplot using fluff (REF XX) and used as input for motif scanning analysis using the *maelstrom* function in Gimme Motifs (Bruse & van Heeringen, 2018). Motifs with in at least one of the clusters a z-score > 3, were selected and visualized with its potential binding transcription factors using an adjusted version of the *Motif2TF* function of scANANSE (Smits et al., 2023).

#### scANANSE analysis on Multiome data

The Seurat implementation of scANANSE (AnanseSeurat, v1.1.0) was used to obtain the important transcription factor regulators from the Multiome dataset (Smits et al., 2023). The resulting differential network for the aCM cluster over vCM cluster, was filtered on the top 20 transcription factors with the highest influence score. Interactions between the nodes with a score > 0.8 were selected from the full cell type-specific networks. Subsequent filtering with the three RA-induced gene sets (described in 7.4.18) from the EB analysis, was used to visualize the regulatory hierarchy within the GRN. The GRN visualizations were generated with Cytoscape v3.9.1 (Shannon et al., 2003). The *ZNF711* network was selected from the aCM full network, keeping only interactions with an interaction score > 0.8 and if they were found as factors of influence in this cell type or known cardiac markers.

#### Validation KOs RNA-seq, lineage marker enrichment and differential analysis

The Seq2Science v0.9.6 pipeline was used to map the RNA-seq reads to the GRCh38 genome (van der Sande et al., 2023). Regularized log transformation was performed with the R package DESeq2 v1.34.0 prior to Principal Component Analysis and heatmap visualization of the gene counts (Love et al., 2014). For each of the day 14 EB cell types from the atlas, the top 200 marker genes were selected per cell type lineage. An average expression level per 200 markers was calculated for each of the bulk RNA-seq samples and corrected for the average expression of the *TRAC* control samples of both conditions, allowing direct comparison between the culture conditions. Significance was tested with a paired Wilcox Sign test for each of the replicates with its matching *TRAC* control sample, which was corrected for multiple testing using the Benjamini & Hochberg method, indicated in the heatmap (p-adjusted < 0.05, < 0.001 and < 0.0001) are based on the highest p-adjusted value per set of replicates. Differential gene expression analysis between *TRAC* control and targeted conditions was performed with DESeq2. The differential gene list was filtered for a log2 fold change > 1 and p-adjusted value < 0.05, and visualized with pheatmap v1.0.12 and EnhancedVolcano v1.12.0. Gene ontology analysis was performed with clusterProfiler v4.2.0 (Wu et al., 2021).

#### Hypergeometric tests on overlap scANANSE predicted targets

The hypergeometric test was performed with the *phyper* function in R. Per knockout and +RA or non-RA condition, the list of differentially expressed genes was used as *successes* and the overlapping genes between these lists and the scANANSE predicted targets of the *ZNF711*, *LHX2* or *RARB*, were considered the *sample successes* in the hypergeometric tests. All genes (16,232 in total) present in both the mutiome scRNA-seq modality and the bulk RNA-seq samples, were considered the full population. For the comparison of ZNF711 targets with the targets of all other regulators in the epicardial, aCM and vCM networks predicted by scANANSE, all interactions per regulator with a probability of 0.8 were kept from the full network. Within each of the networks the Jaccard Index (the targets overlap divided by the union of the two target-sets minus the overlap) was used to calculate the amount of overlap between the predicted ZNF711 targets and the target sets of each other regulator in the network and enrichment was tested with a hypergeometric test, with all genes (26,128 in total) expressed in the multiome dataset as full population. The p-values from the hypergeometric tests were corrected for multiple testing using the Benjamini & Hochberg method. Calculated p-values smaller than 1e-100 (including zero values) were reported as p < 1e-100.

## DATA AVAILABILITY

All NGS data is available on GEO with the temporal scRNA-seq data at GSE263193, the multiome day 8 EB samples at GSE263326 and the RNA-seq data of the KO samples at GSE263325. Published data was obtained from GSE106118 (Cui et al., 2019) or as filtered matrices published on the author’s website (Asp, 2021; Asp et al., 2019). The single-cell temporal atlas object for visualization can be downloaded through Zenodo via https://doi.org/10.5281/zenodo.10932845. The per-cluster pseudobulk scATAC-seq profiles can be viewed in the Genome Browser at https://genome.ucsc.edu/s/rebecza/scRoadmap_CardiacDiffs_hg38.

## CODE AVAILABILITY

All used software is mentioned in the METHODS, the custom R and Python scripts used for the analyses, as well as instructions on how to visualize the single-cell atlas data in an interactive user interface are available on Github at https://github.com/Rebecza/scRoadmap_CardiacDiffs.

## SUPPLEMENTARY DATA

**Supplementary Table S1:** Motif activity and scANANSE analysis on the day 8 embryoid body multiome.

**Supplementary Table S2:** Differential gene expression and gene ontology analysis on *ZNF711*, *LHX2* and *ZNF503* knockouts.

**Supplementary Table S3:** Input for scANANSE and comparison with DE-genes from *ZNF711* knockout differentiations to atrial and ventricular lineages.

## AUTHOR CONTRIBUTIONS

R.R.S. conceived experiments, generated libraries for all Next Generation Sequencing (NGS) methods used in the manuscript, led the computational analysis and wrote the manuscript. C.C.F. conceived experiments, performed all cell culture work used in the manuscript, harvested and sorted the cells, generated the CRISPR-Cas9 knockout lines and wrote the manuscript. M.B. generated bulk RNA-sequencing libraries and performed the sequencing for all NGS libraries used in the manuscript. V.S. conceived experiments. R.P. conceived experiments, supervised the experiments involving stem cell differentiation, wrote the manuscript and generated funding for the work. G.J.C.V. conceived experiments, supervised computational genomics analysis, wrote the manuscript and generated funding for the work.

## ACKNOWLEDGEMENTS

This work was funded by a ZonMw TOP grant (project number 91217061) to R.P. and G.J.C.V.

## DISCLOSURE AND COMPETING INTEREST STATEMENT

R.P. is a cofounder of Pluriomics (Ncardia) and River BioMedics BV. The other authors declare no competing interests.

## Notes

### Summary of Updates

The manuscript has been revised based on reviewer reports. Additional analyses have been performed and incorporated in the manuscript. Textual mistakes have been corrected, and the clarity of the text has been improved based on the feedback received.

