## Supplementary Figures for "Cardiac differentiation roadmap for analysis of plasticity and balanced lineage commitment"

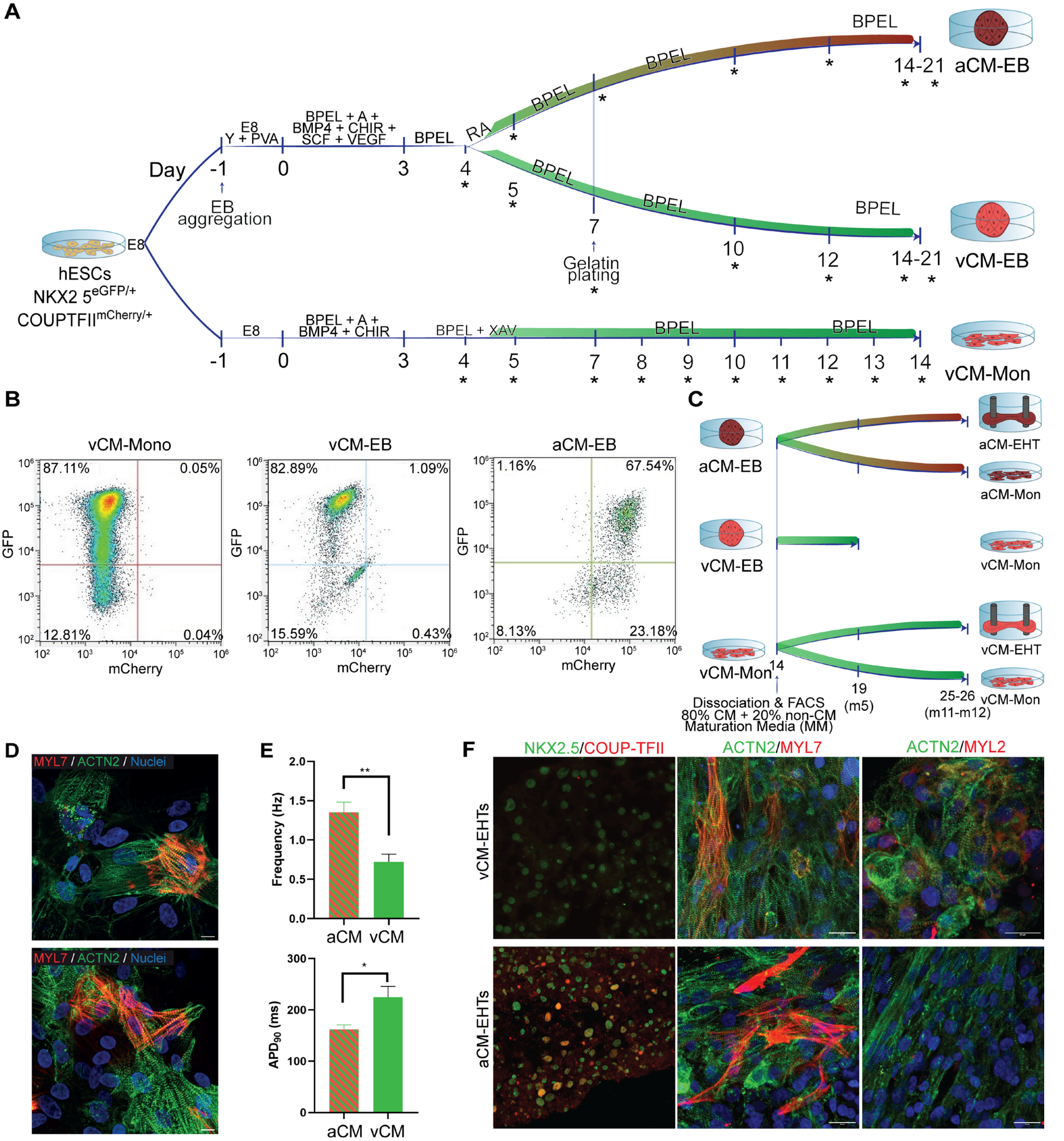


**Supplementary Figure S1. Atrial and ventricular differentiation protocols and CM characterization**.

(A) Detailed overview of the differentiation protocols from hPSC towards atrial or ventricular embryonic bodies (aCM-EBs, vCM-EBs, respectively) or ventricular monolayers (vCM-Mon). The stars indicate the timepoints at which single cells were harvested per protocol for single-cell RNA-sequencing. Abbreviations: E8 = Essential 8 media; BPEL = Bovine Serum Albumin Polyvinylalchohol Essential Lipids media; A = Activin A; BMP4 = Bone Morphogenetic protein 4; CHIR = Chir99021; SCF = Stem Cell Factor; VEGF = Vascular Endothelial Growth Factor; RA = retinoic acid analog; XAV = xav939; m = day in maturation medium.

(B) Representative flow cytometry plots depicting the differentiation efficiency of the resultant cardiomyocytes at day 14 from each of the differentiation protocols used in (A), using the reporter line NKX-2.5$^{eGFP+/-}$-COUP-TFII$^{mCherry+/-}$. Atrial CMs are both GFP and mCherry positive while ventricular CMs only GFP positive.

(C) Schematic overview of the protocol used at day 14 to generate EHTs and monolayer controls per culture type. Note from vCM-EB, only monolayers were generated since EHTs from vCM-Mon were more efficient to generate. All indicated timepoints were harvested as single cells for single-cell RNA-sequencing. m5-m12 = day 5-12 in maturation medium (day 14-26; EHT and monolayer only).

(D) Additional immunohistochemistry of general cardiomyocyte markers MYL7 and ACTN2 in day 21 vCM-EBs (top) and aCM-EBs (bottom). Scale bar = 20 µm.

(E) Frequency (top) and action potential duration at 90% of repolarization (APD90) (bottom) of aCMs and vCMs EBs at day 14, determined by FluoVolt voltage sensitive dye. Data are mean ± s.e.m. Two-tailed Student's t-test. *$p < 0.05$; **$p < 0.01$. (n = 6 EBs from 3 batches of differentiation).

(F) Representative confocal images of ventricular and atrial EHTs (aCM-EHTs, vCM-EHTs, respectively) stained for NKX-2.5, COUP-TFII, ACTN2, MYL7 and MYL2 confirming their lineage identity. Scale bar = 20 µm.

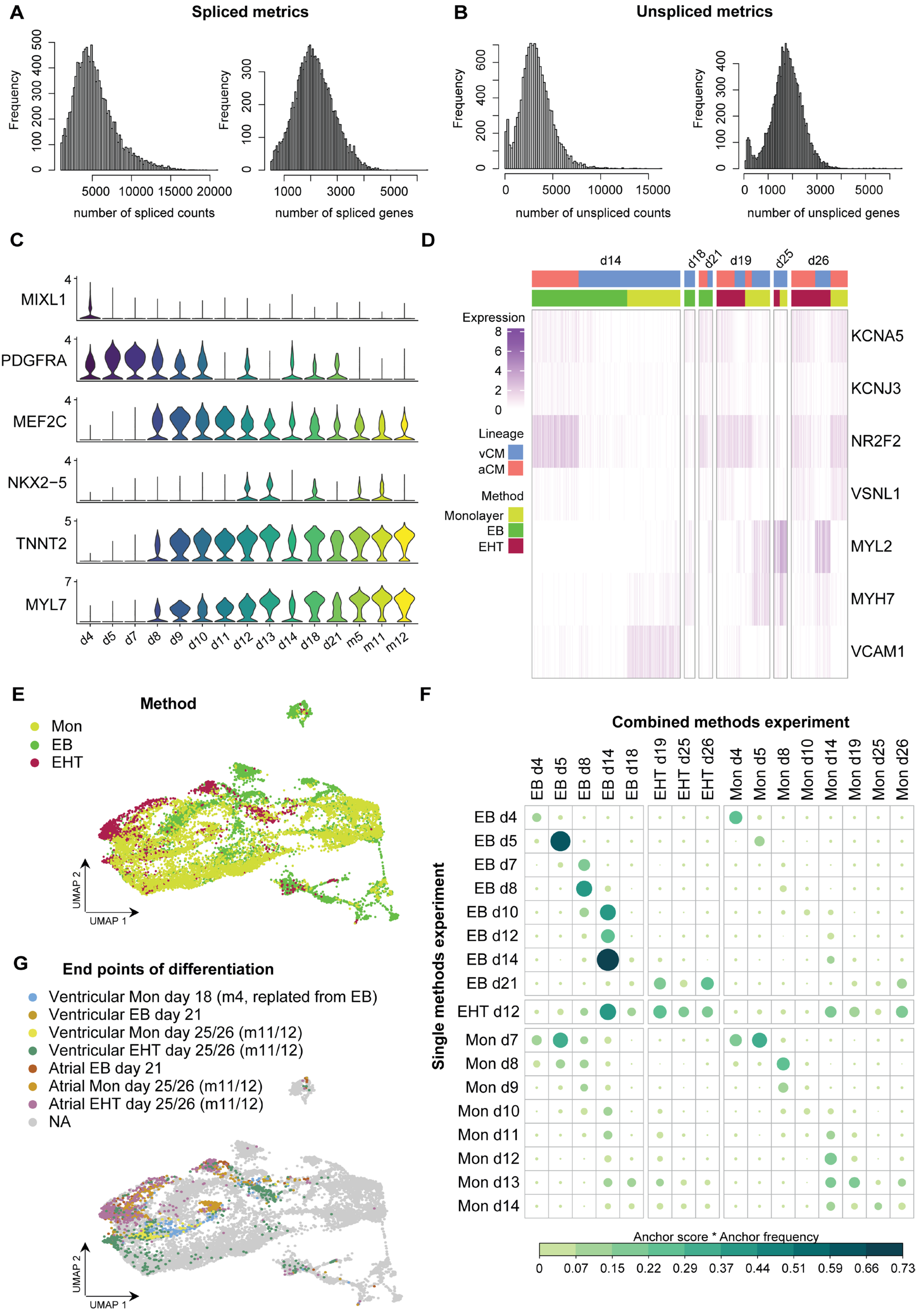

A
Spliced metrics
Frequency
number of spliced counts
number of spliced genes
B
Unspliced metrics
number of unspliced counts
number of unspliced genes
C
MIXL1
PDGFRA
MEF2C
NKX2−5
TNNT2
MYL7
d4 d5 d7 d8 d9 d10 d11 d12 d13 d14 d18 d21 m5 m11 m12
D
d14 d18 d21 d19 d25 d26
Expression
Lineage
vCM
aCM
Method
Monolayer
EB
EHT
KCNA5
KCNJ3
NR2F2
VSNL1
MYL2
MYH7
VCAM1
E
Method
Mon
EB
EHT
UMAP 1
UMAP 2
F
Combined methods experiment
EB d4 EB d5 EB d8 EB d14 EB d18 EHT d19 EHT d25 EHT d26 Mon d4 Mon d5 Mon d8 Mon d10 Mon d14 Mon d19 Mon d25 Mon d26
Single methods experiment
EB d4 EB d5 EB d7 EB d8 EB d10 EB d12 EB d14 EB d21 EHT d12 Mon d7 Mon d8 Mon d9 Mon d10 Mon d11 Mon d12 Mon d13 Mon d14
Anchor score * Anchor frequency
0 0.07 0.15 0.22 0.29 0.37 0.44 0.51 0.59 0.66 0.73
G
End points of differentiation
Ventricular Mon day 18 (m4, replated from EB)
Ventricular EB day 21
Ventricular Mon day 25/26 (m11/12)
Ventricular EHT day 25/26 (m11/12)
Atrial EB day 21
Atrial Mon day 25/26 (m11/12)
Atrial EHT day 25/26 (m11/12)
NA
UMAP 1
UMAP 2

**Supplementary Figure S2. Single-cell RNA-sequencing dataset.**

(A-B) Histogram showing the frequency of cells counted in the dataset after quality filtering with (A) spliced (exonic) or (B) unspliced (intronic) reads, summarized as read counts (left) and detected genes (right) per cell (x-axis).

(C) Combined expression of mesoderm (*MIXL1*, *PDGFRA*), cardiac progenitor (*MEF2C*, *NKX2-5*) and cardiomyocyte (*MYL7*, *TNNT2*) marker genes across the full dataset per time point.

(D) Heatmap representation of atrial and ventricular markers for evaluation of lineage identity in atrial (salmon-red) and ventricular (blue) cultures from day 14 to day 26. Yellow-green, green and carmine-red indicate monolayer, embryoid body and engineered heart tissue respectively.

(E) UMAP representation of the integrated dataset as shown in Fig. 1C-D, cells labelled for the method (top).

(F) Dotplot visualizing the frequency and score of mutual nearest neighbors (integration anchors) between the culture experiments, columns show the samples in one experiment where all methods were cultured in parallel, the rows indicated separate experiments for each of the culture methods. Size and color of the dots represents the product of anchor score and anchor frequency.

(G) UMAP representation of the integrated dataset as shown in Fig. 1C-D, cells labelled for the end points of differentiation for each of the different methods (full protocol of differentiation is shown in Figure 1A, Supplementary Fig. S1A,C). m4-m11-m12 = day 4-11-12 in maturation medium (day 18-25-26; EHT and monolayer only).

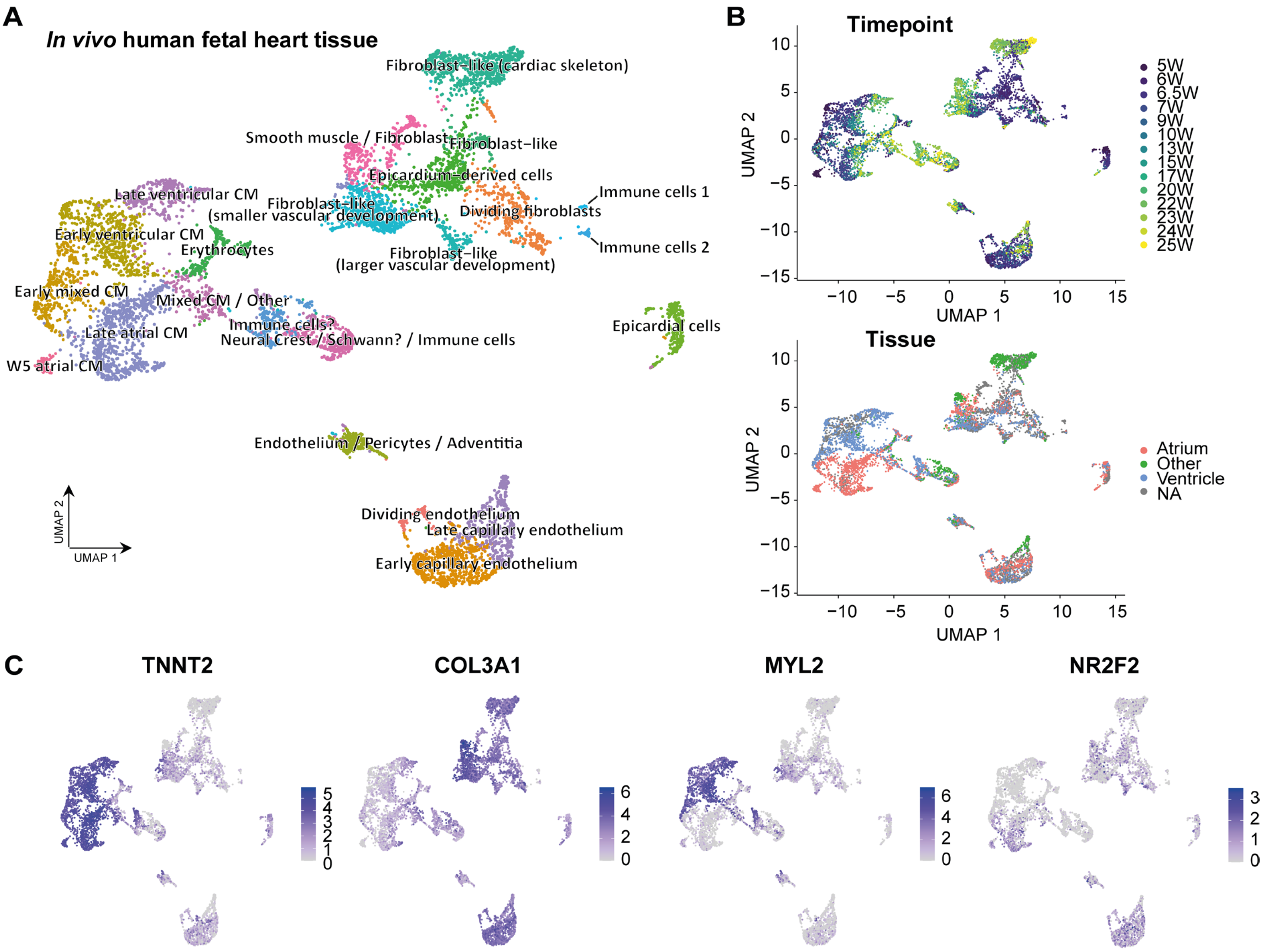


**Supplementary Figure S3. Cell types of the human fetal heart.**

(A) Integrated scRNA-seq datasets of human fetal heart tissue from Cui et al. (2019) and Asp et al. (2018), labelled with extended annotation based on the cell types annotated in the Asp et al. dataset.

(B) UMAP representation as shown in (B) labelled with time point (top) and tissue (bottom) of harvest where this information was provided (w, weeks after conception).

(C) Gene expression of known markers *TNNT2* (general cardiac muscle marker), *MYL2* (ventricular cardiomyocyte marker), *NR2F2* (atrial cardiomyocyte marker), and *COL3A1*.

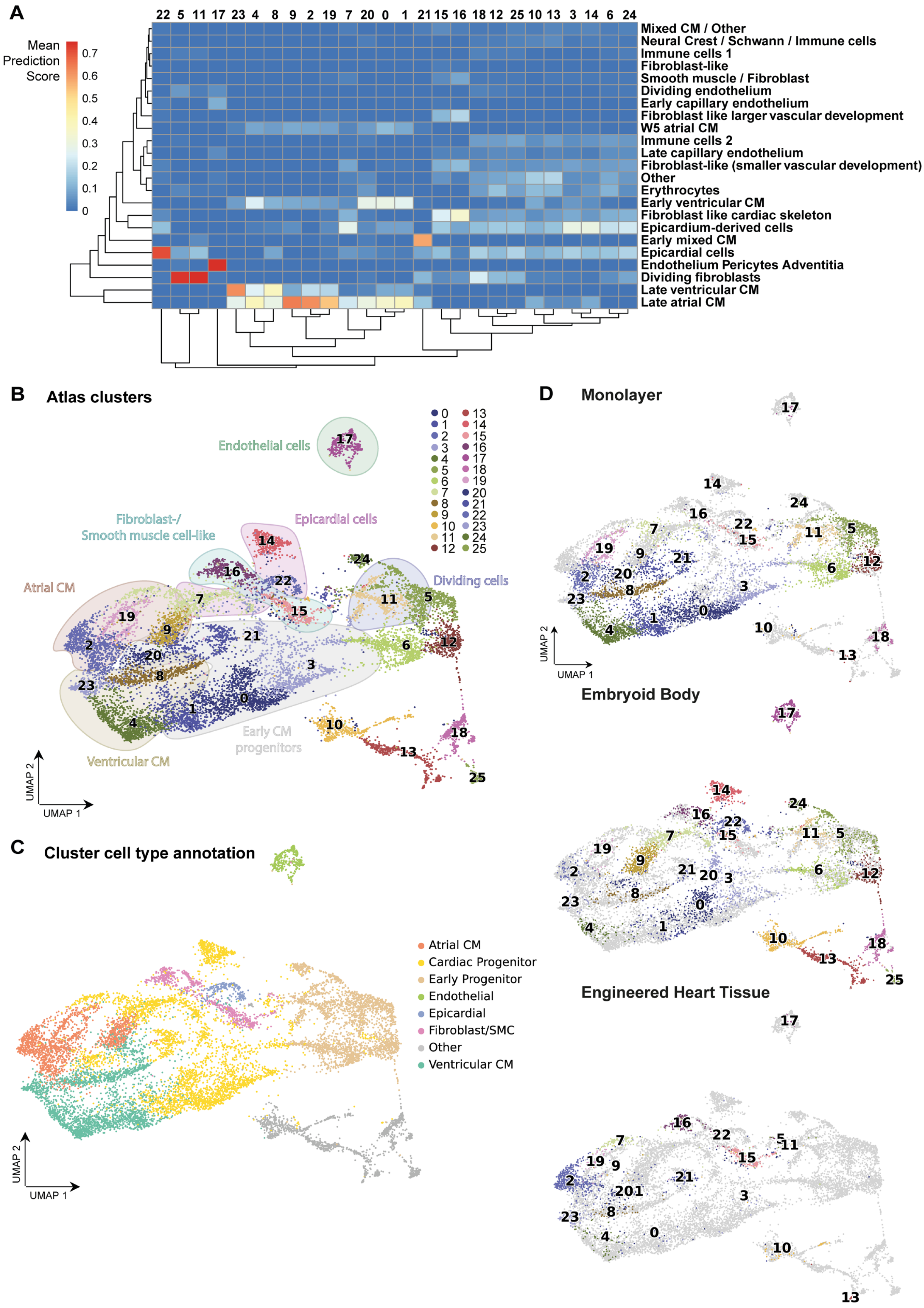


**Supplementary Figure S4. Annotating the temporal single-cell atlas.**

(A) Heatmap with predicted fetal cell types from Figure S3 (rows) for each of the atlas clusters from Figure 1C and S3A (columns). Multiple late time point clusters (day 14 and older) from the ventricular and atrial lineage, show most similarity to in vivo vCM and aCM clusters, respectively. Cluster 22 matched best with epicardial and cluster 17 with endothelial cells. Early clusters were similar to proliferating fibroblasts (cluster 5 and 11).

(B) UMAP representation of the atlas with cluster labelling, colored stains show the annotation process based on the scoring in (A).

(C) UMAP representation of the atlas with the final cell state annotation.

(D) UMAP representation of the full atlas, with coloring of the clusters as shown in (B), separated per culture method. Cells in light grey fall outside of the culture method of focus.

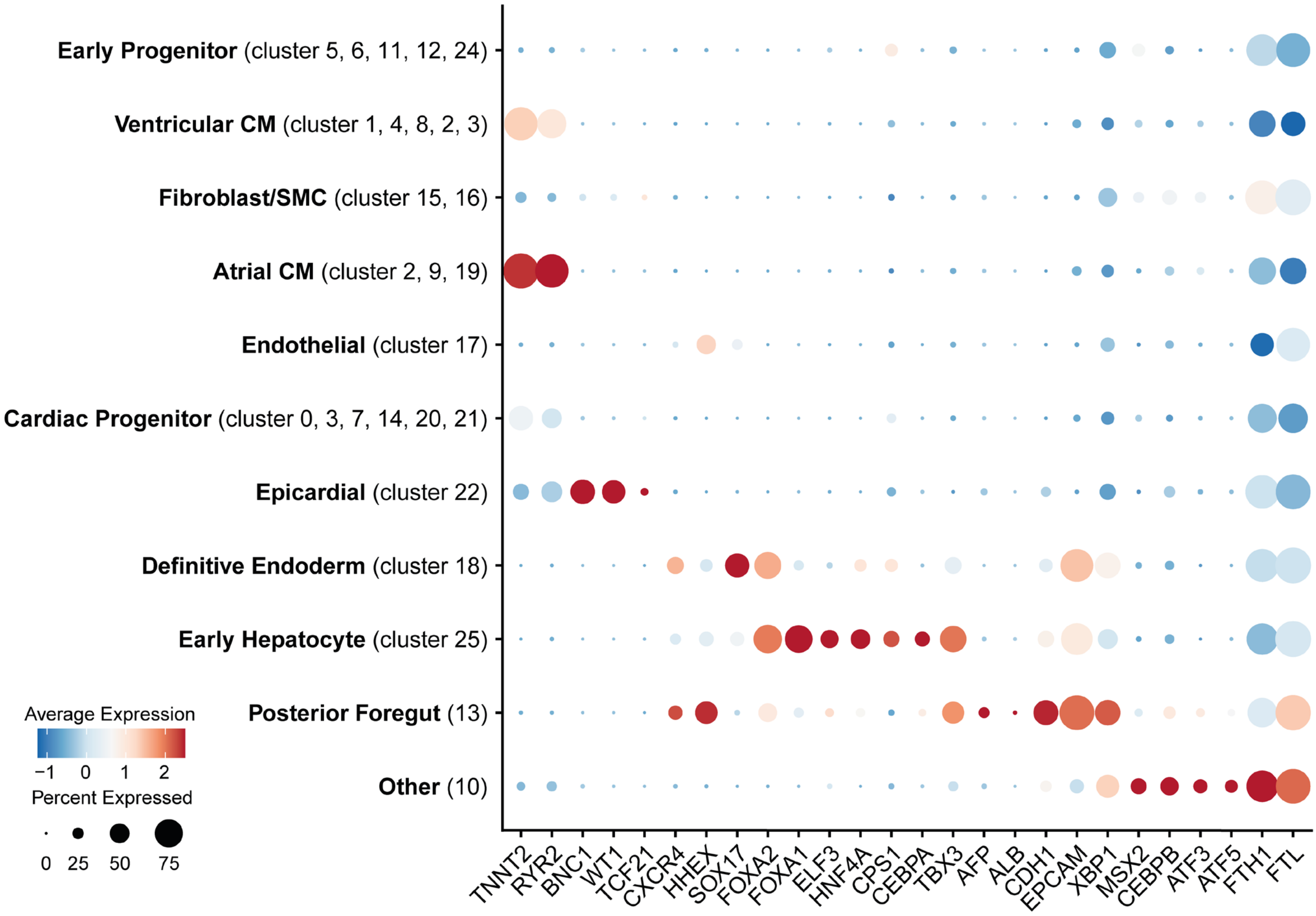


**Supplementary Figure S5.**

Dotplot showing marker genes for the different clusters: general cardiomyocyte *TNNT2*, *RYR2*; epicardial *BNX1*, *WT1*, *TCF21*; and markers for definitive endoderm (*SOX17*, *CXCR4* (or *CD184*), *FOXA1*, *FOXA2*), early hepatocyte (*FOXA1*, *ELF3*, *HNF4A*, *CPS1*, *CEBPA*, *TBX3*, *AFP*, *ALB*), (posterior) foregut (*ELF3*, *HHEX*, *CDH1*, *EPCAM*), and marker genes found in the Other cluster (*MSX2*, *CEBPB*, *ATF3*, *ATF5*, *FTH1*, *FTL*) (Ang et al., 2018; Scheibner et al., 2021).

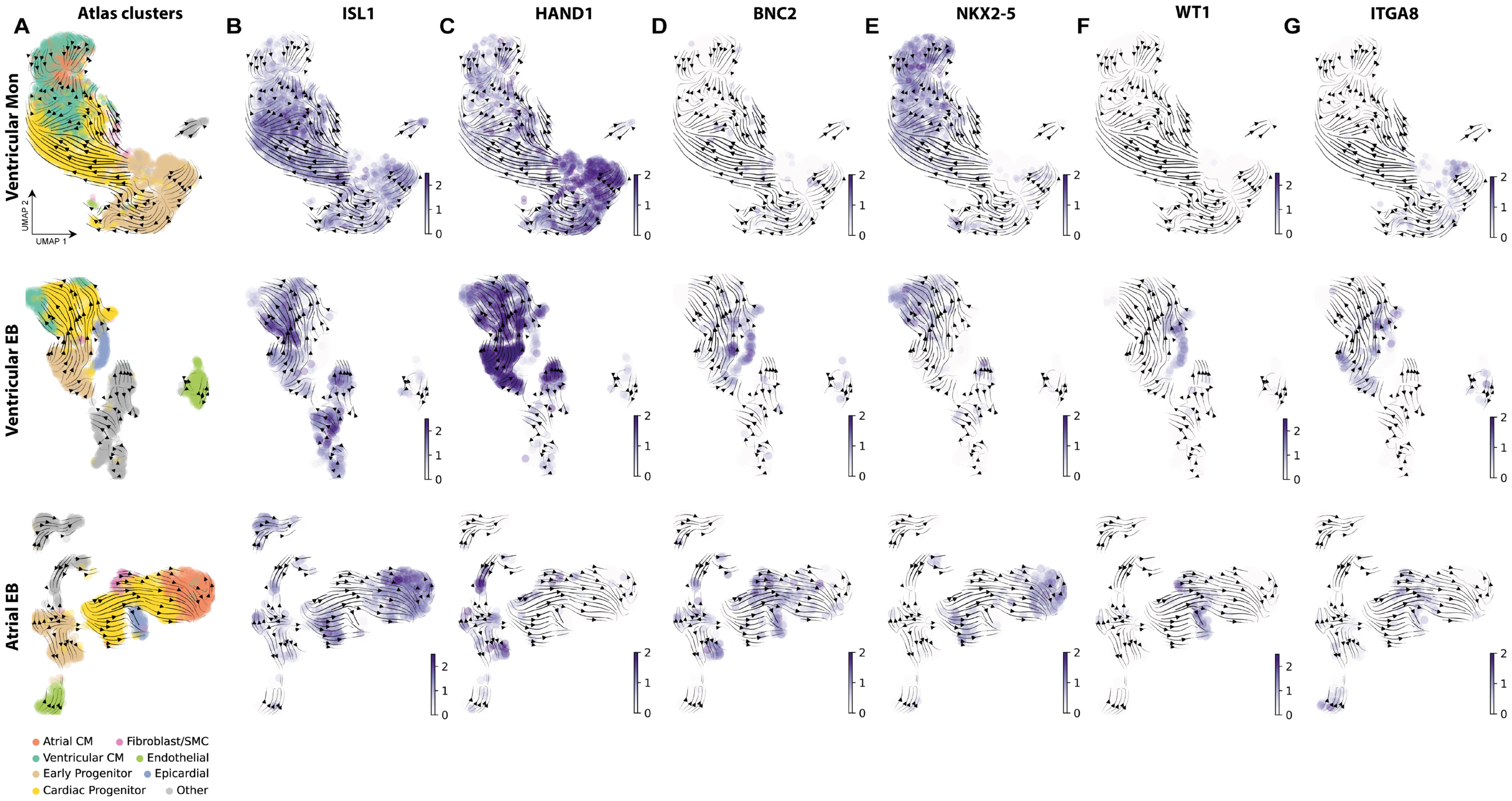


**Supplementary Figure S6: Early trajectories with SHF and early CM markers ISL1 and NXK2-5 and (pre-)epicardial marker expression.**

Per culture protocol UMAP representation with ventricular monolayer culture (top), ventricular embryoid body (EB) (middle) and atrial EB (bottom) labeled for cell types (A) and expression levels of SHF marker *ISL1* (B), epicardial marker *WT1* (C) and genes expressed in pre-epicardial cells *HAND1* (D), *BNC2* (E), *NKX2-5* (F), as well as the new epicardial surface marker *ITGA8* (G) (Zawada et al., 2023).

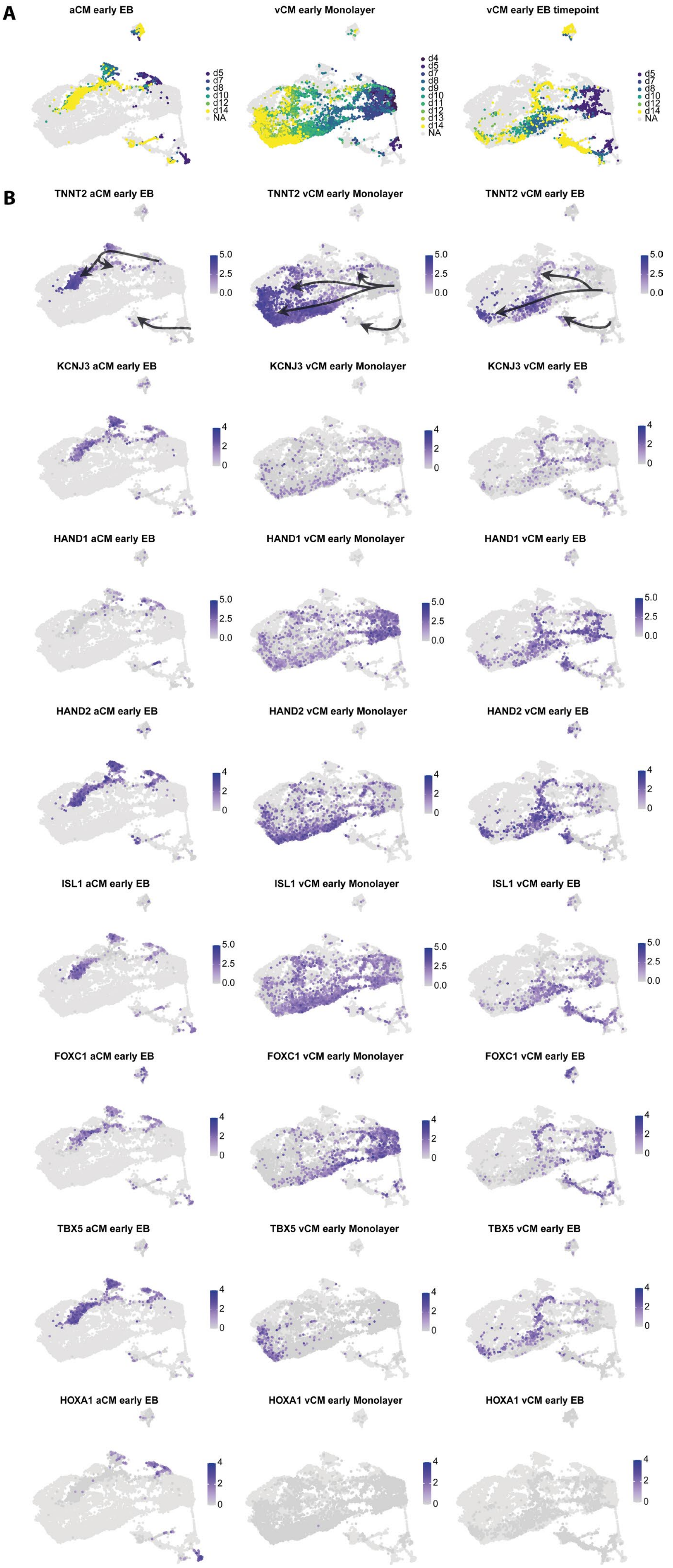
A
aCM early EB
vCM early Monolayer
vCM early EB timepoint
d5
d7
d8
d10
d12
d14
NA
d4
d9
d11
d13
B
TNNT2 aCM early EB
TNNT2 vCM early Monolayer
TNNT2 vCM early EB
KCNJ3 aCM early EB
KCNJ3 vCM early Monolayer
KCNJ3 vCM early EB
HAND1 aCM early EB
HAND1 vCM early Monolayer
HAND1 vCM early EB
HAND2 aCM early EB
HAND2 vCM early Monolayer
HAND2 vCM early EB
ISL1 aCM early EB
ISL1 vCM early Monolayer
ISL1 vCM early EB
FOXC1 aCM early EB
FOXC1 vCM early Monolayer
FOXC1 vCM early EB
TBX5 aCM early EB
TBX5 vCM early Monolayer
TBX5 vCM early EB
HOXA1 aCM early EB
HOXA1 vCM early Monolayer
HOXA1 vCM early EB
5.0
2.5
0.0
4
2
0

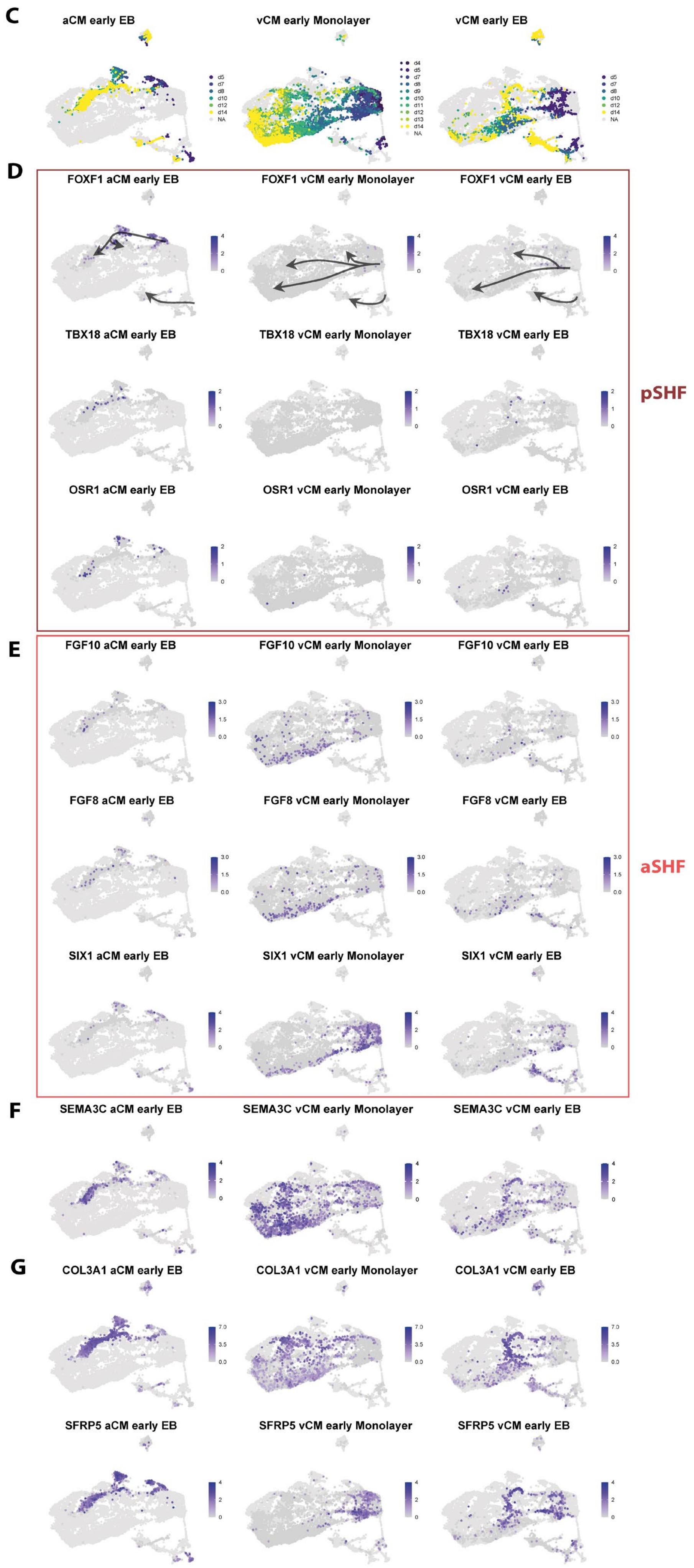

C
aCM early EB
vCM early Monolayer
vCM early EB
d4
d5
d7
d8
d9
d10
d11
d12
d13
d14
NA
D
FOXF1 aCM early EB
FOXF1 vCM early Monolayer
FOXF1 vCM early EB
TBX18 aCM early EB
TBX18 vCM early Monolayer
TBX18 vCM early EB
OSR1 aCM early EB
OSR1 vCM early Monolayer
OSR1 vCM early EB
pSHF
E
FGF10 aCM early EB
FGF10 vCM early Monolayer
FGF10 vCM early EB
FGF8 aCM early EB
FGF8 vCM early Monolayer
FGF8 vCM early EB
SIX1 aCM early EB
SIX1 vCM early Monolayer
SIX1 vCM early EB
aSHF
F
SEMA3C aCM early EB
SEMA3C vCM early Monolayer
SEMA3C vCM early EB
G
COL3A1 aCM early EB
COL3A1 vCM early Monolayer
COL3A1 vCM early EB
SFRP5 aCM early EB
SFRP5 vCM early Monolayer
SFRP5 vCM early EB

**Supplementary Figure S7: Early trajectory expression patterns.**

(A) Atlas UMAP representation labelled with time point of harvest per culture protocol, cells from other protocols in the atlas are shown in light grey (NA).

(B) Expression levels of genetic markers for the cardiomyocytes (*TNNT2*), atrial CM (*KCNJ3*), first heart field (*HAND1*), second heart field (*HAND2*, *ISL1*, *FOXC1*, *TBX5*).

(C) Same representation as in (A). UMAP labelled with time point of harvest per culture protocol, cells from other protocols in the atlas are shown in light grey (NA).

(D) Expression levels of genetic markers for the posterior second heart field (pSHF).

(E) Expression levels of genetic markers for the anterior second heart field (aSHF).

(F) Expression levels of *SFRP5* and *COL3A1*, which showed elevated co-expression in the non-CM branches of the UMAP, where *SFRP5* was downregulated and to a lower extent *COL3A1* in the ventricular cardiomyocyte branch (middle and right), whereas both remain upregulated in atrial CM differentiation (left).

(G) Expression levels of *SEMA3C* marker of aSHF-derived ventricular cardiomyocytes.

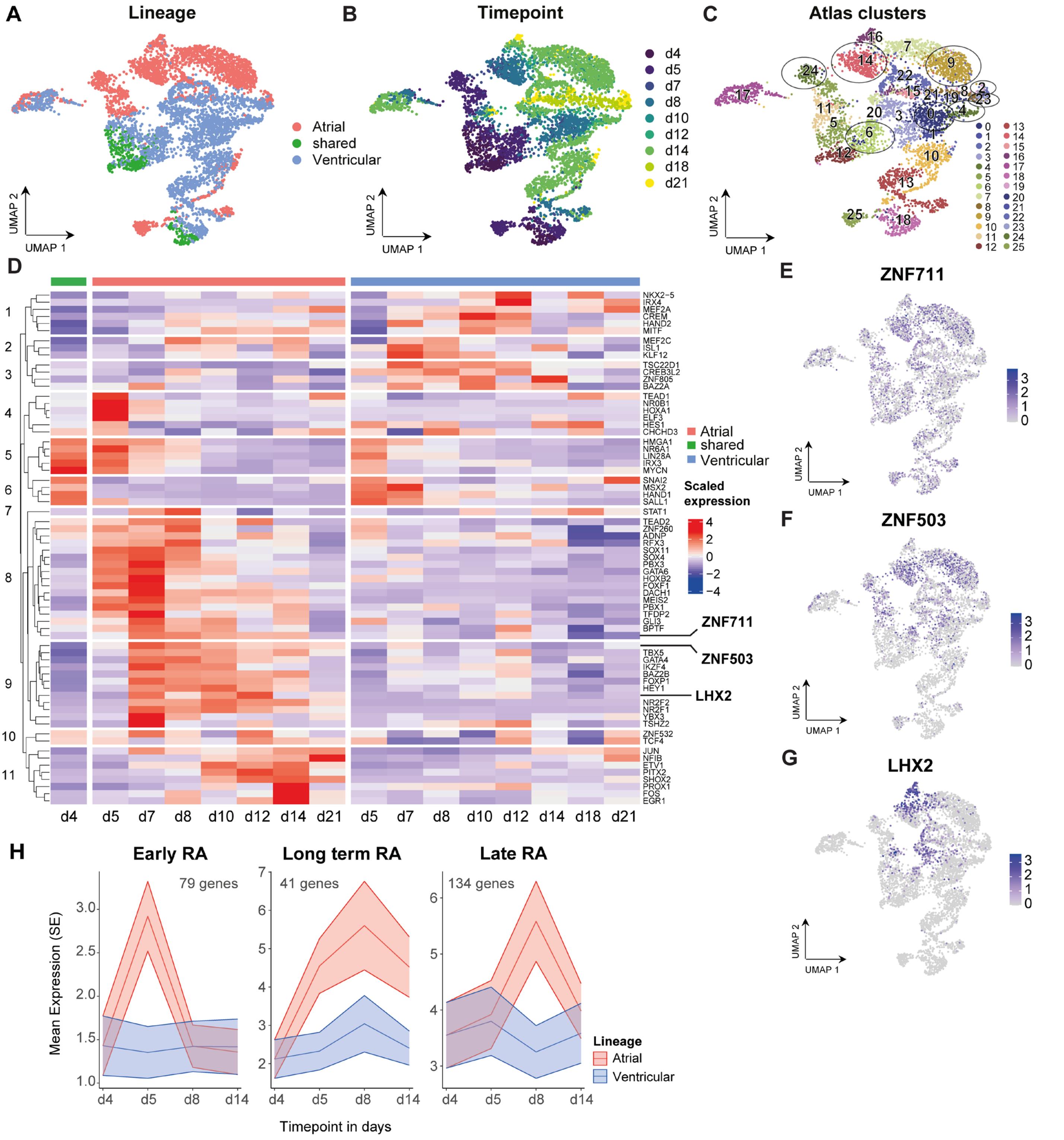


**Supplementary Figure S8. Transcription factor analysis in the embryoid body (EB) atlas subset.**

(A-B) UMAP representation of EB subset of the atlas, cells labelled with the lineage they were cultured towards (A) and the time point the cells were harvested (B).

(C) UMAP with circles outlining the differential gene analysis between the lineages performed on atlas clusters in trajectory to aCM and vCM. Comparisons were performed between cluster 24 and 6, cluster 14 and 0, cluster 9 and 4, and between cluster 2 and 23 were made.

(D) Heatmap showing the results of differential expression analysis between early and late cardiomyocyte clusters, outlined in (B), across the two lineages within the EB dataset.

(E-G) Highlighted hits of interest from (B), *ZNF711* (E), *ZNF503* (F) and *LHX2* (G) visualized with their expression levels over the UMAP representation of the EB subset.

(H) Average expression levels over time per gene set which was found differentially upregulated in atrial versus ventricular culture, subdivided into an early RA-responsive (left), a long-term RA-responsive (middle) and late RA-responsive subset (right) (*see Methods*). d = day of culture, day 4 is shared between the lineages and day 5 the first day after administration of RA in the atrial culture.

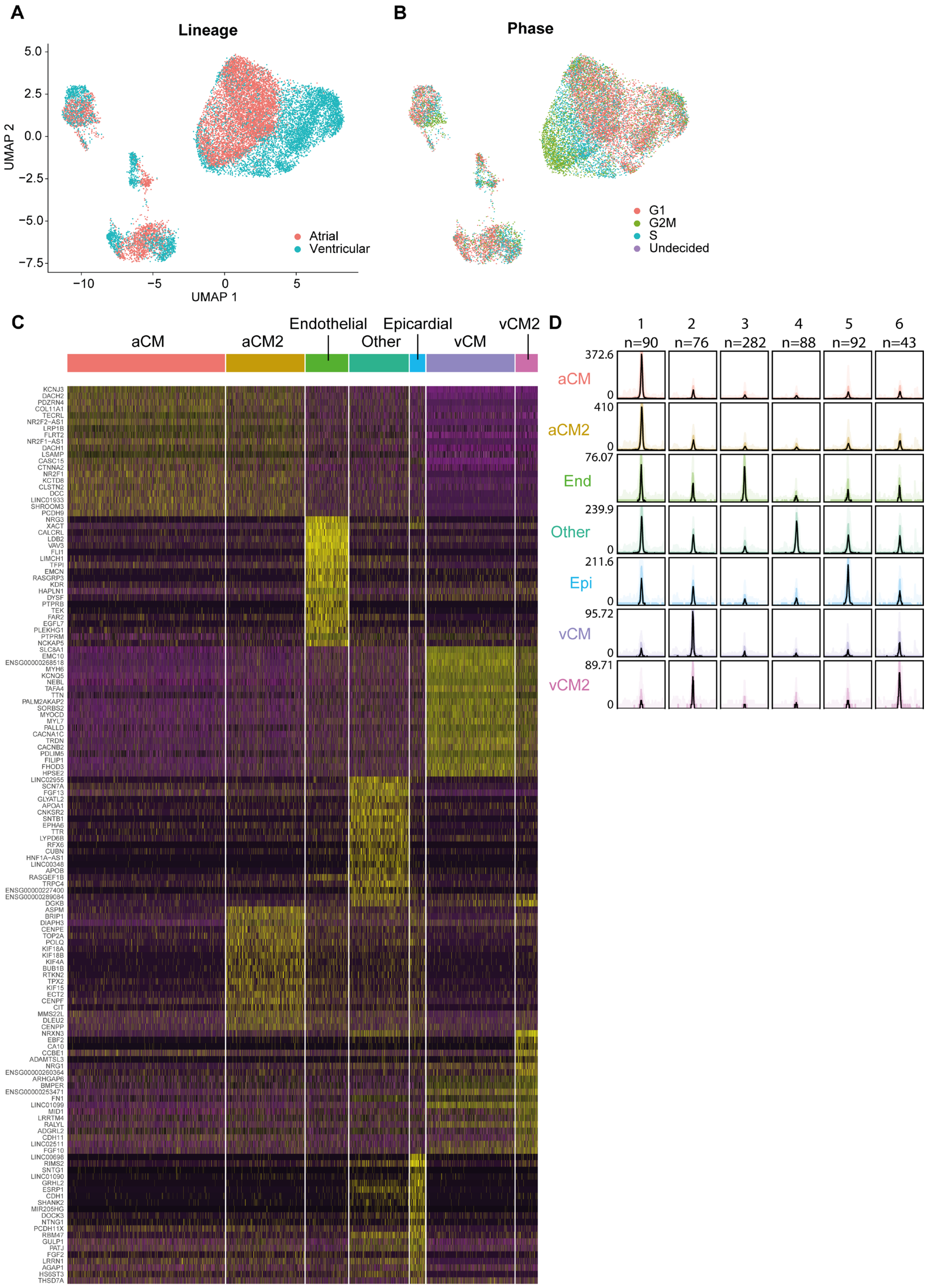


**Supplementary Figure S9. Cluster identification and patterns of gene expression and chromatin accessibility in the multiome dataset.**

(A-B) UMAP representation from Figure 3c with labels of the lineage they were cultured towards (A) and the predicted cell cycle phase (B).

(C) Heatmap visualizing expression levels of top 20 genetic markers, called per cell type.

(D) Chromatin accessibility patterns per cell type in the multiome dataset with a bandplot showing a graph illustration of differentially accessible peaks per cluster. These peaks were used for motif analysis (Supplementary Table S1A-B).

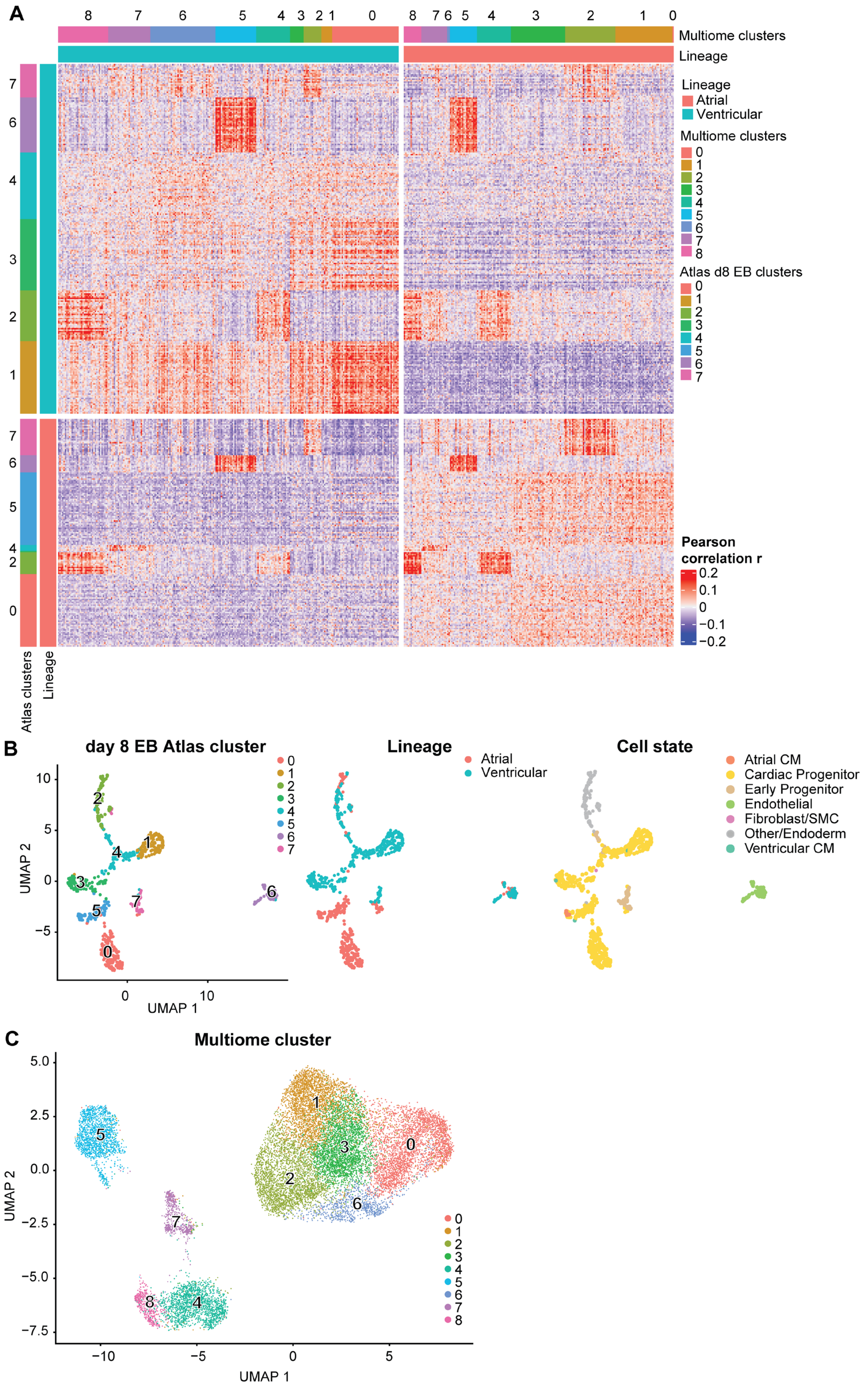
A
Multiome clusters
Lineage
Lineage
Atrial
Ventricular
Multiome clusters
Atlas d8 EB clusters
Pearson correlation r
Atlas clusters
B
day 8 EB Atlas cluster
Lineage
Atrial
Ventricular
Cell state
Atrial CM
Cardiac Progenitor
Early Progenitor
Endothelial
Fibroblast/SMC
Other/Endoderm
Ventricular CM
UMAP 1
UMAP 2
C
Multiome cluster
UMAP 1
UMAP 2

**Supplementary Figure S10: Correlations between day 8 EB subset of the temporal Atlas and day 8 multiome data.**

(A) Correlation heatmap with random subsets of 1,000 cells per cluster in the multiome dataset for the columns and all cells per cluster of the atlas dataset in the rows, additionally separated per lineage.

(B) UMAP shown with clusters called for the d8 EB subset of the atlas (left), labelled for lineage origin (middle) and labeled for cell types as annotated in the atlas (right).

(C) UMAP representation of the multiome dataset labelled with the cluster numbers.

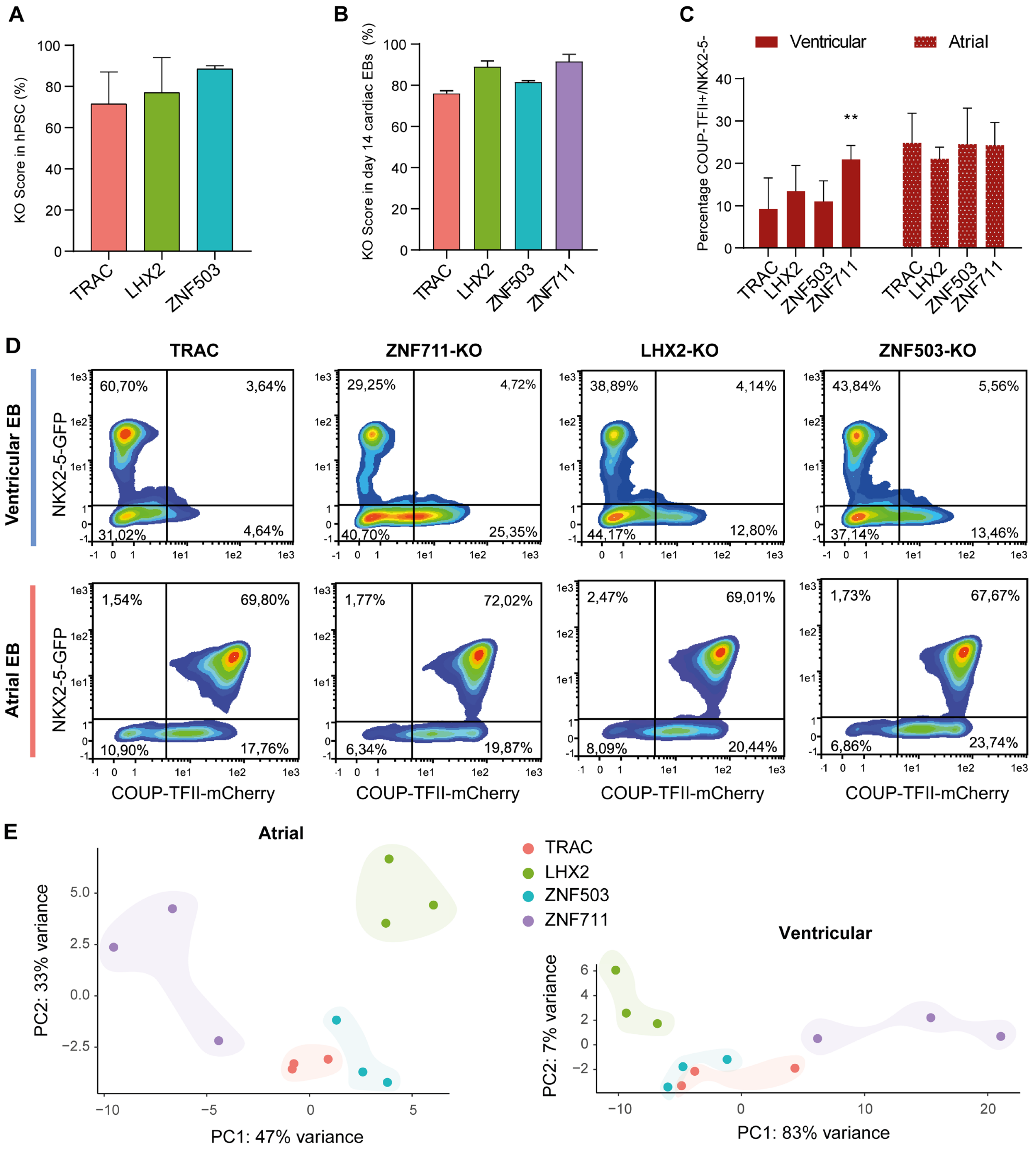


**Supplementary Fig S11 Transcription factor knockout experiment read-out.**

(A) Knockout (KO) efficiency per target of hPSCs at the start of differentiation, *TRAC* is used as a CRISPR control. Data are mean ± s.e.m; ordinary One-way ANOVA with Tukey's multiple comparisons test. (n=2 independent hPSC cultures per KO target). ZNF711-KO missing due to sample loss.

(B) KO efficiency per target after differentiation of each KO-hPSC culture to atrial or ventricular CMs, measured at day 14 of embryoid body (EB) culture. Data are mean ± s.e.m; ordinary One-way ANOVA with Tukey's multiple comparisons test. (n=4 per target, resulting from two independent EB differentiations per lineage).

(C) Percentages of COUP-TFII (NR2F2)-positive and NKX2-5-negative cells per KO target measured by flow cytometry in non-RA-treated (ventricular, left red bars) and +RA-treated (atrial, red, white-dotted bars) conditions. Data are mean ± s.e.m; ordinary Two-way ANOVA with Tukey's multiple comparisons test. **p < 0.001. (n=3 independent differentiations per culture type).

(D) Representative flow cytometry results from panel (C), showing on the y-axis the signal measured for NKX2-5-GFP and on the x-axis the signal measured for COUP-TFII-mCherry, represented with a density plot for each of the control conditions (TRAC) and KO conditions, cultured with ventricular (non-RA, top row) or atrial (+RA, bottom row) conditions. Atrial CMs are both NKX2-5-GFP and COUP-TFII-mCherry positive while ventricular CMs only GFP positive. Non-CMs are either NKX2-5-GFP negative or both NKX2-5-GFP negative and COUP-TFII-mCherry positive.

(E) PCA on RNA-seq of KO target samples obtained after 14 days of EB differentiation separate for atrial (plus RA; top) and the ventricular lineage (no RA; bottom).

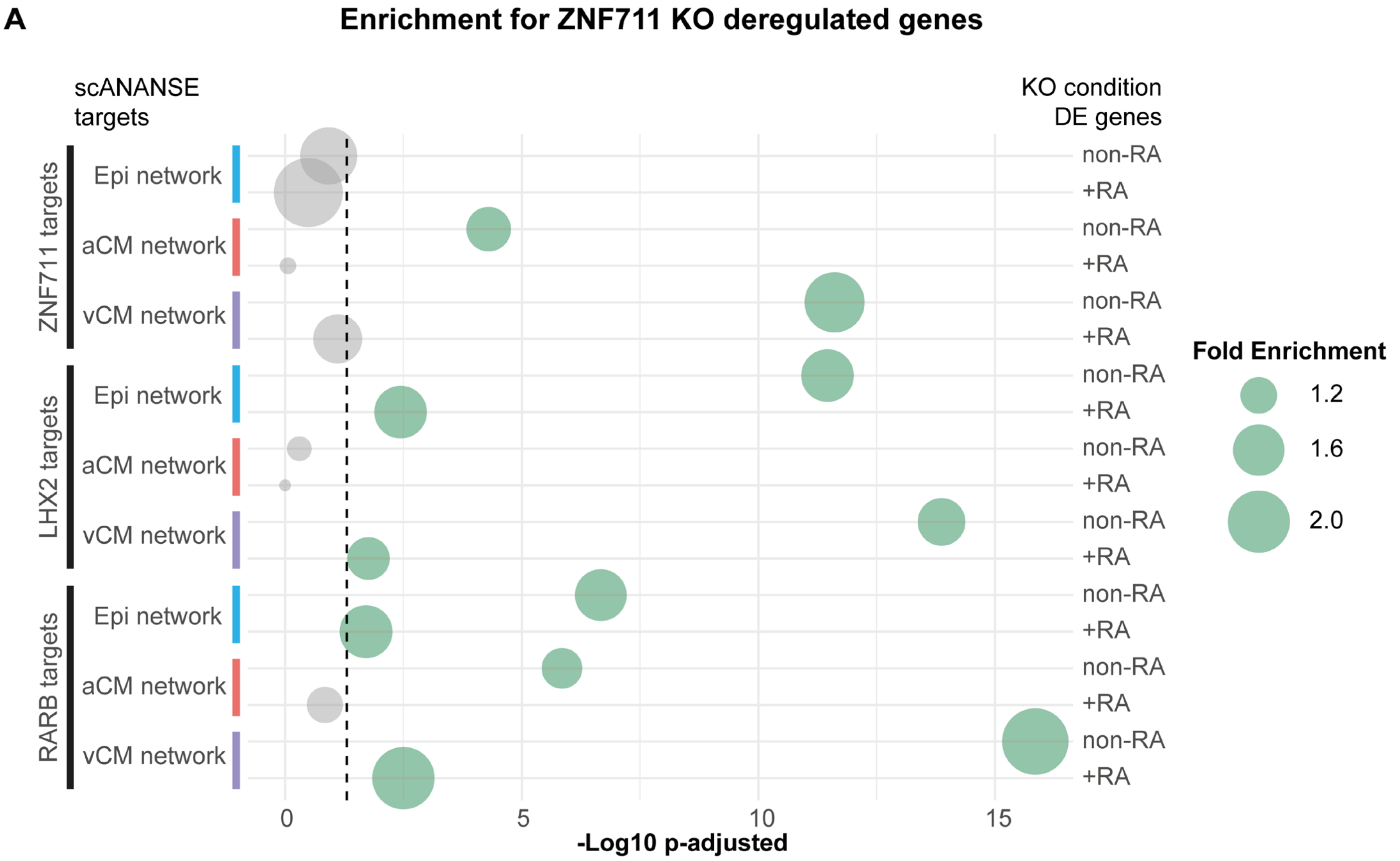


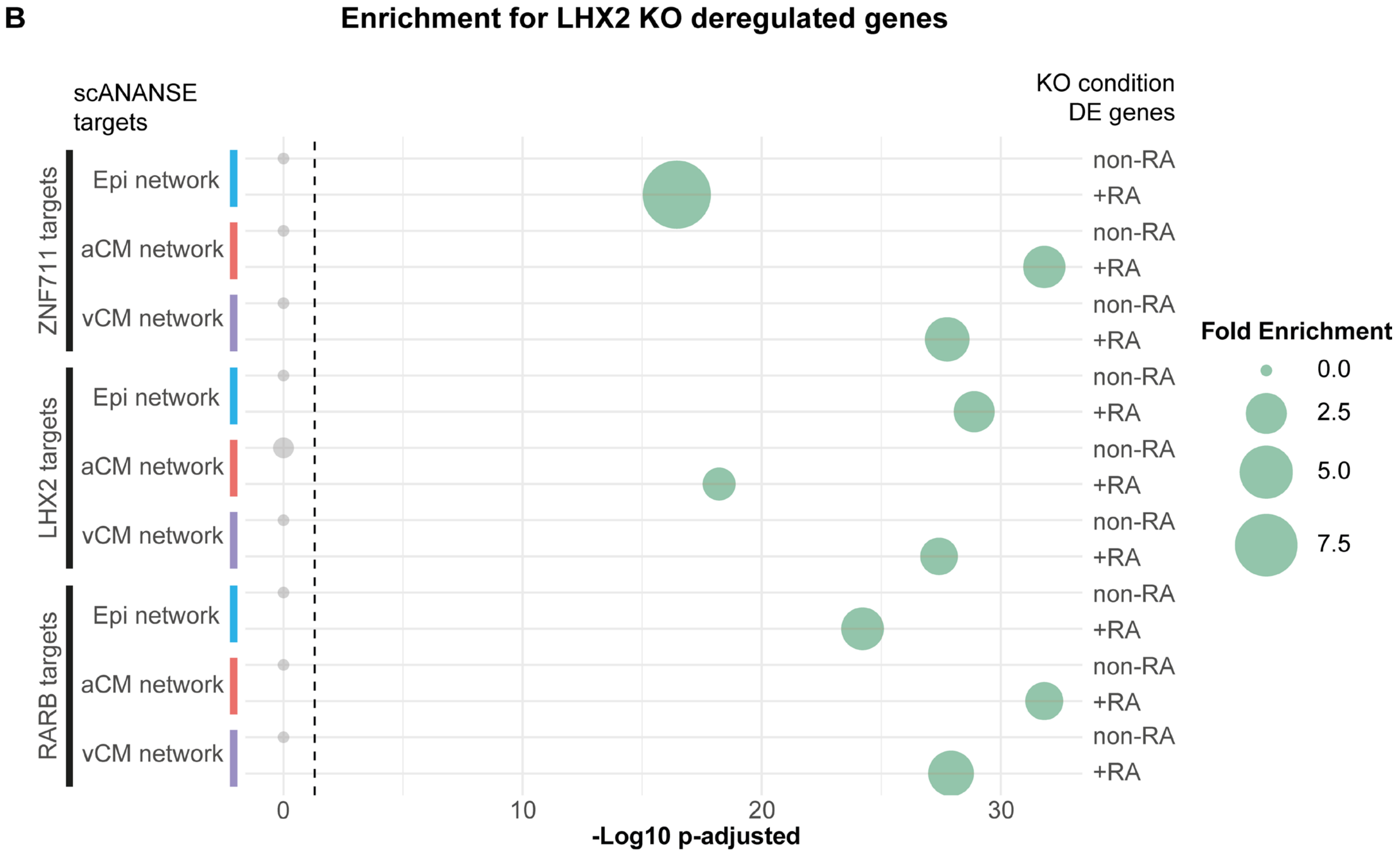


**Supplementary Figure S12: All enrichment results of K.O. targets in predicted ANANSE networks.**

(A-B) Enrichment and significance (hypergeometric test, multiple testing-corrected) of the ZNF711 KO targets (A) or LHX2 KO targets (B), overlapping with the predicted scANANSE targets for ZNF711, LHX2 and RARB, for each of the cell type networks. Indicated on the left of the bubble plot are the regulators for which the predicted targets were selected and from which cell type-specific scANANSE network. On the right of the plot, the conditions are indicated from which the *ZNF711*-KO differential genes were selected, non-RA (ventricular) or +RA (atrial). The dashed line indicates the threshold for a significant p-adjusted value of 0.05.

Epi = epicardial cells, aCM = atrial cardiomyocyte, vCM = ventricular cardiomyocyte.
